# Effects of Aging, Fitness, and Cerebrovascular Status on White Matter Microstructural Health

**DOI:** 10.1101/2024.08.04.606520

**Authors:** Grace M. Clements, Paul Camacho, Daniel C. Bowie, Kathy A. Low, Bradley P. Sutton, Gabriele Gratton, Monica Fabiani

## Abstract

White matter (WM) microstructural health declines with increasing age, with evidence suggesting that improved cardiorespiratory fitness (CRF) may mitigate this decline. Specifically, higher fit older adults tend to show preserved WM microstructural integrity compared to their lower fit counterparts. However, the extent to which fitness and aging *independently* impact WM integrity across the adult lifespan is still an open question, as is the extent to which cerebrovascular health mediates these relationships. In a large sample (N = 125, aged 25-72), we assessed the impact of age and fitness on fractional anisotropy (FA, derived using diffusion weighted imaging, DWI) and probed the mediating role of cerebrovascular health (derived using diffuse optical tomography of the cerebral arterial pulse, pulse-DOT) in these relationships. After orthogonalizing age and fitness and computing a PCA on whole brain WM regions, we found several WM regions impacted by age that were independent from the regions impacted by fitness (hindbrain areas, including brainstem and cerebellar tracts), whereas other areas showed interactive effects of age and fitness (midline areas, including fornix and corpus callosum). Critically, cerebrovascular health mediated *both* relationships suggesting that vascular health plays a linking role between age, fitness, and brain health. Secondarily, we assessed potential sex differences in these relationships and found that, although females and males generally showed the same age-related FA declines, males exhibited somewhat steeper declines than females. Together, these results suggest that age and fitness impact specific WM regions and highlight the mediating role of cerebrovascular health in maintaining WM health across adulthood.

## 1. Introduction

Aging is associated with many brain changes, including a reduction in total gray and white matter brain volume and consequent enlargement of the ventricles and subarachnoid space (e.g., Narvacan et al., 2017). White matter volumes (i.e., macrostructure) are typically stable until ≈ age 60, after which a precipitous decline begins (Ge et al., 2002; Raz et al., 2005; Walhovd et al., 2005), whereas diffusion-tensor measures of white matter, which are sensitive to the magnitude and direction of water diffusion, precede macrostructural changes and therefore have increased sensitivity to detect changes across the lifespan (Sullivan & Pfefferbaum, 2006; Zahr et al., 2010). In the last two decades, cardiorespiratory fitness (CRF), which can be improved via regular aerobic exercise, has been shown to mitigate the impact of brain aging in later life (for a review see Bowie, Clements, et al., 2021). Indeed, evidence suggests that higher levels of CRF are associated with preserved gray and white matter volumes (Colcombe et al., 2006) in both cortical (Fletcher et al., 2016; Gordon et al., 2008) and subcortical areas (Erickson et al., 2011; McAuley et al., 2011; Niemann et al., 2014).

As a modifiable lifestyle factor, CRF provides an attractive means to reduce the impact of aging on the brain. However, fitness may not globally and equally preserve all brain regions. As such, a more nuanced approach may be needed to understand the possible regional effects of CRF. Indeed, Fletcher et al. (2016) reported that increased fitness and advanced age selectively influenced distinct gray matter volumes, cortically and subcortically. A potential explanation for these differential effects of aging and fitness on brain tissue integrity is through the health of the cerebrovasculature, a vast network of arteries, arterioles and capillaries that envelops the brain, providing it with life-sustaining oxygen and glucose and enabling clearance of unwanted substances (Zimmerman et al., 2021). In midlife, as part of the normal aging process, arteries begin to harden (arteriosclerosis; Bowie, Clements et al., 2021; Izzo & Shykoff, 2001). This reduces cerebrovascular function (Fabiani et al., 2022) and may impact brain tissue health if certain areas are not sufficiently perfused. In the cross-sectional study reported here, we investigate the unique contributions of aging and fitness to the integrity of the white matter across the adult lifespan, as well as sex differences and the potential mediating role of cerebrovascular health in these relationships.

Since the mid-2000s, diffusion-weighted magnetic resonance imaging (dwMRI) has been leveraged to study how the microstructural integrity of the white matter varies across the lifespan, by quantifying the diffusion of water molecules in brain tissues (Assaf & Pasternak, 2008; Bennett et al., 2010; Gunning-Dixon et al., 2009; Sullivan & Pfefferbaum, 2006). Differently from white matter macrostructural volumes, which tend to be fairly preserved until the sixties or seventies (Ge et al., 2002; Raz et al., 2005; Walhovd et al., 2005), white matter *microstructural* changes become apparent in midlife (in the thirties or forties; Beck et al., 2021; Sullivan & Pfefferbaum, 2006; Westlye et al., 2010; Zahr et al., 2010). This increased variability in midlife makes the microstructure better suited to study early age-related changes in white matter; in particular, one dwMRI metric often assessed in aging is fractional anisotropy (FA). FA is a normalized measure of the variance of water molecule diffusion across the 3 primary directions within a voxel, with larger values associated with greater cellular organization within the measured tissue (Wozniak & Lim, 2006). With increasing age, FA in the white matter tends to decline globally (Hsu et al., 2008; Hugenschmidt et al., 2008; Kochunov et al., 2012; Westlye et al., 2010), with consistent reports of decreased FA in the corpus callosum (Sullivan et al., 2001; Salat et al., 2005), among other regions. As such, we predict that FA will decrease across the lifespan, but that changes will not be uniformly distributed across all white matter.

Considerable evidence from observational and interventional studies, as well as cross-sectional and longitudinal studies, supports the positive impact of fitness on white matter integrity in aging (cross-sectional: Hayes et al., 2015; Oberlin et al., 2016; Tarumi et al., 2021b; Tian et al., 2014; Longitudinal: Burzynska et al., 2017; Clark et al., 2019; Mendez Colmenares et al., 2021; Voss et al., 2013). An excellent recent meta-analysis without publication bias (Maleki et al., 2022) found strong evidence for the positive impact of physical activity, CRF, and exercise on FA in the corpus callosum and anterior limb of the internal capsule, indicating preserved white matter integrity with increased fitness in these regions. However, of the extant dwMRI observational, cross-sectional studies that specifically assess the relationship between CRF and white matter integrity, none assesses the entire adult lifespan and instead tend to focus either on older adults only (N.F. Johnson et al., 2012; Liu et al., 2012; Marks et al., 2011; Oberlin et al., 2016; Tian et al., 2014), younger adults only (Herting et al., 2014; Zhu et al., 2015), or compare extreme groups of younger and older adults (Hayes et al., 2015). To our knowledge, only two studies (d’Arbeloff et al., 2021; Tarumi et al., 2021a) include the middle age range, a critical age at which white matter degradation begins to occur, and in these two studies the middle-age group was examined in isolation rather than in the context of the entire adult lifespan. We address this gap here by including middle-aged individuals in the context of a broad age range (ages 21-71).

Some changes in cardiovascular and cerebrovascular function begin to emerge in middle age, including the onset of arteriosclerosis (the loss of arterial elasticity; Najjar et al., 2005; Sun, 2015). Peripherally, elevated pulse pressure (the difference between systolic and diastolic blood pressure) can be used as an approximate index of increasing arteriosclerosis (Dart & Kingwell, 2001; Franklin, 2005; Franklin et al., 1997). Here we use diffuse optical tomography of the cerebral arterial pulse (pulse-DOT, Fabiani et al., 2014; Gratton et al., 2017; Tan et al., 2016; 2017; 2019; Chiarelli et al., 2017; Bowie et al., submitted; Kong et al., 2020) to assess *cerebral* arteriosclerosis. Pulse-DOT metrics of arterial elasticity have been reliably related to age, CRF, brain volumetric atrophy, and brain function (Chiarelli et al., 2017; Fabiani et al., 2014; Kong et al., 2019; Tan et al., 2017, 2019). Given that the onset of vascular dysfunction typically begins in middle age (AlGhatrif et al., 2013; Vasan et al., 2002), it is imperative to include this group when assessing white matter health, which is especially impacted by the hypoperfusion caused by arteriosclerosis (Badji et al., 2019; Tan et al., 2019; Zimmerman et al., 2021), among other vascular factors. Indeed, white matter macrostructural lesion volume is linked to vascular damage (Tarumi et al., 2015) and controlling arterial stiffness has been shown to slow lesion progression (Prins & Scheltens, 2015), indicating that vascular impacts are particularly evident in the white matter compared to gray matter and that altering the vasculature can mitigate these negative effects.

Here we start with the hypothesis that aging is associated with reduced white matter integrity, but that at least in certain age groups and for certain brain regions, enhanced fitness diminishes some of these reductions. Aging is also associated with reduced cerebrovascular function, but enhanced fitness ameliorates or at least staves off some of these changes as well (Aatola et al., 2014). Note that, although we predict that higher levels of CRF will be related to more preserved white matter integrity, it is not necessarily the case that the regions impacted by fitness are perfectly overlapping with those impacted by aging. Indeed, it may be that fitness exerts both direct and indirect effects on the white matter. For example, the white matter regions that are implicated in exercise may be better preserved in highly fit individuals (total effect)—but it is also known that engaging in more physical activity may also lead to a reduction/slowing in arteriosclerosis/arterial stiffness, thereby preserving white matter health in those areas for which vascular effects are more important (indirect effect). The regions indirectly affected by vascular health may not overlap with those affected by complementary exercise-induced mechanisms, including increases in neurotrophic factors such as brain-derived neurotrophic factor (BDNF), vascular endothelial growth factor (VEGF), and insulin-like growth factor 1 (IGF-1), which were not collected in the present study (Voss et al., 2011). Therefore, a **mediating** effect would be supported by an indirect effect of *fitness* on white matter integrity via *cerebrovascular health* or an indirect effect of *age* on white matter integrity via *cerebrovascular health*. A **moderating** effect would be supported by a significant interaction between *age* and *fitness* on white matter health.

With increasing fitness, there tends to be a concomitant *increase* in arterial elasticity. With increasing age, there tends to be a concomitant *decrease* in arterial elasticity. Given these antagonistic effects, we hypothesize that, a) cerebral measures of arterial health may mediate the relationship between age and white matter integrity and/or fitness and white matter integrity and, b) if fitness has a protective effect against age-related declines in white matter microstructure, we may find an interaction between fitness and aging on certain white matter tracts, thereby indicating moderation. Lastly, given that biological sex impacts the regulation of vascular tone (Thompson & Khalil, 2003) partially due to the anti-inflammatory and vasoprotective effects of estradiol (an estrogen) in females (Gilligan et al., 1994; Hurn et al., 1995; see also Meyer et al., 2006; Reckelhoff, 2018) resulting in differences in arterial elasticity before and after menopause (Rossi et al., 2011), we sought to identify potential differences between females and males in our sample by analyzing their data separately.

In summary, our primary aim was to assess the relative impact of aging and fitness on white matter microstructural integrity. Our secondary aims were to assess the potentially mediating role of cerebrovascular health therein, as well as potential sex differences in these relationships. To preview the results, in a cross-sectional, observational adult sample of over 100 individuals, we found several white matter regions impacted by age that were independent from the regions impacted by fitness, whereas other areas showed interactive effects of aging and fitness. Measures of cerebrovascular health mediated the relationships between aging or fitness and white matter microstructure. Females and males did not have equivalent white matter integrity in all brain regions, and although both generally showed the same age-related white matter integrity declines, males tended to decline at a faster rate than females. Together, these findings support the impact of fitness on white matter microstructural integrity across the adult lifespan as well as between sexes and paint a dynamic picture of the complex interactions between these factors.

## 2. Method

All code used in this project can be found at https://osf.io/j9psq/

### 2.1 Participants and Procedure

Unpublished data from 127 participants collected in our lab were analyzed in this study (mean age = 53.47 yrs, range = 25-71, 35% male). The data were pooled across two large studies that used nearly identical data collection protocols for the variables of interest (73 participants from Study 1, age range 50-70; and 54 participants from Study 2, age range 25-71)^1^. All participants were native English speakers and reported themselves to be in good health with normal hearing and normal or corrected-to-normal vision, and free from medications that may directly affect the central nervous system. All participants signed informed consent and all procedures were approved by the Institutional Review Board of the University of Illinois Urbana-Champaign. Specific criteria for inclusion for this report were a usable dwMRI scan and an estimated CRF measure (eCRF). One participant had an extreme eCRF metric due to a very high BMI (more than 5 standard deviations above the mean), and a second had extreme head motion in the scanner (framewise displacement more than 4 standard deviations above the mean) and were thus excluded from analysis, rendering the final sample size 125. In this sample, 113 (90.4%) participants self-reported as White, 9 (7.2%) as Asian, 2(1.6%), as Black or African American and 1 (0.8%) as multiracial. In terms of ethnicity, 5 (4%) self-identified as Hispanic/Latino and 120 (96%) as not Hispanic/Latino.

### 2.2 Data Acquisition

#### 2.2.1 Cardiorespiratory Fitness Assessment

We used Jurca et al. (2005) estimation of cardiorespiratory fitness (eCRF), which uses 5 values (age, sex, BMI, resting heart rate, and activity score) to compute a metabolic equivalent (MET) of VO2_max_. Many studies have used this metric and shown that it is highly correlated (≈0.7) with VO2_max_ (Jurca et al., 2005; Stamatakis et al., 2013). eCRF has been validated in older adults (Mailey et al., 2010; McAuley et al., 2011).^2^ Participants’ age, sex, body mass index, resting heart rate and a self-reported activity score are weighted and then linearly combined with a constant to estimate CRF, as follows:

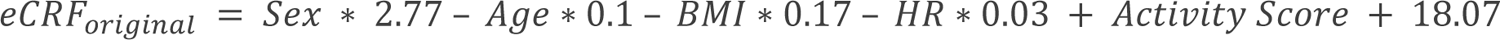

In our sample, resting heart rate was calculated as the average of 3 measurements taken at rest during 3 different laboratory visits. eCRF defined this way retains the biological reality that, on average, males have greater VO2_max_ than females because of their larger bodies and lungs. To control for these known systematic differences that would not be expected to reflect real variations in fitness or vascular health, we effectively removed sex from the eCRF equation by omitting the sex term from the calculations and then adding 2.77/2 to all subjects. In this way, instead of the sex term being 0 for females and 2.77 for men, all participants have a constant 1.385 included in their eCRF estimations, allowing for the direct comparison of males and females:

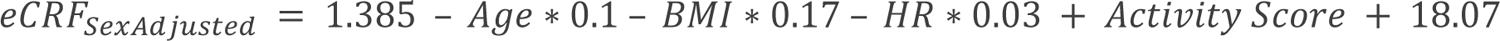

Please note that, although VO2_max_, assessed with a graded exercise test, is considered the gold standard for determining CRF, we used eCRF in this study for the following reasons. First, as mentioned, several groups (Jurca et al., 2005; Mailey et al., 2010; McAuley et al., 2011) have shown that eCRF is an accurate method to quantify CRF across a wide age range like the one used in this study, thereby increasing the validity of our CRF estimates. Second, in our current sample only participants in Study 1 completed a graded exercise test, so we would severely limit our sample size by focusing exclusively on VO2_max_. Third, the graded exercise test is a challenging experience, and some older participants’ health places limitations on their ability to complete the test. As a result, many older adults in our sample who *did* complete the test *did not* reach their peak VO2, yielding graded exercise test measures for this subset of participants less reliable (Abdelkarim et al., 2023). Finally, a demonstration that eCRF relates to white matter microstructural integrity across the adult lifespan is missing from the literature.

#### 2.2.2 Structural MRI Acquisition

Each participant underwent an MRI session, which included acquisition of a T1-weighted and a diffusion weighted sequence on a Siemens Prisma 3T scanner with a 20-channel head coil. T1-weighted images were acquired with a magnetization prepared gradient-echo (MPRAGE) sequence with the following parameters: TR/TE/TI = 2400 ms / 2.31 ms / 1060 ms; flip angle = 8°; acquisition matrix = 240 × 256 × 180; iPAT factor of 2; 0.8 mm isotropic voxels, with the prescan normalize option. Slices were acquired in the sagittal plane.

dwMRI data were acquired with a diffusion weighted sequence with associated reverse phase-encoding directions echo planar imaging field maps and an interleaved multiband spin echo planar imaging (EPI) sequence with a multiband acceleration factor of 4 (Auerbach et al., 2013; Setsompop et al., 2012). Sequence parameters were as follows: TR/TE/flip angle: 2500 ms/90 ms/90°, FOV: 230 × 230 mm, 2.5mm isotropic scan. We obtained one volume of b = 0 and diffusion weighted data along 64 directions at both b1 = 1000 and b2 = 2000 s/mm^2^.

In Study 1 the images acquired at each diffusion weighting (b-value) were sequential; that is, 64 volumes at b = 1000 s/mm^2^ were collected and then 64 b = 2000 s/mm^2^ volumes were acquired. In Study 2, the volumes acquired with b = 1000 and b=2000 s/mm^2^ were interleaved/alternating. Global FA values were compared in age-matched participants from both studies using Welch’s two sample unequal variance t-tests, *t*(25.34) = -0.812, *p* = 0.425. Because we found that the FA values were not affected by this manipulation, the scans from the two studies can be combined.

#### 2.2.3 Diffuse Optical Imaging Data Acquisition

Cerebral arterial elasticity (pulse-DOT) data were obtained during an 8–10-minute resting-state optical imaging session, in which seated participants fixated on a cross at the center of a screen. Optical data were acquired with 12 synchronized multichannel frequency-domain oximeters (ISS Imagent, Champaign, IL) equipped with a total of 128 laser diodes (64 emitting light at 690 nm and 64 at 830 nm) and 48 photomultiplier tubes. Time-division multiplexing was employed so that each detector picked up light from 16 different sources at different times within a multiplexing cycle at a sampling rate of 39.0624 Hz. The light was transmitted to the scalp by using single-optic fibers (0.4 mm core) and from the scalp back to the photomultiplier tubes by using fiber bundles (3 mm diameter). The fibers were held in place using semirigid, custom-built helmets, fitted to the participants based on their head circumference.

After the helmet was set up, the locations of the optodes were marked digitally to improve spatial accuracy during later data processing, including anatomical coregistration with each participant’s structural MRI (T1-weighted scan). Fiducial markers were placed on each participant’s left and right preauricular points and the nasion. These fiducial points, optode locations, and other scalp locations were digitized with a Polhemus FastTrak 3D digitizer (accuracy: 0.8 mm; Colchester, VT) by using a recording stylus and three head-mounted receivers, which allowed for small movements of the head in between measurements. Optode locations and structural MRI data were then coregistered using fiducial alignment and/or surface fitting as described by Chiarelli et al. (2015).

Concurrently, an electrocardiogram (EKG) was recorded using Brain-Vision recorder software and a Brain Products V-Amp 16 integrated amplifier system (Brain Products, Germany). Specifically, lead 1 of the EKG (left wrist referenced to right wrist) was recorded with a sampling rate of 500 Hz and a band-pass filter of 0.1-250 Hz. The exact timing of each R-wave peak was determined by searching for peak points exceeding a voltage threshold (scaled for each participant) and dismissing any peak points outside of the normal range of inter-beat intervals. The identification of each peak was verified by visual inspection, and false detections, usually misidentifying the T-wave as the R-wave, were eliminated. Concurrent acquisition of EKG and optical pulse data allowed us to time-lock the optical pulse data to the R-wave of the EKG, thereby ensuring that the same pulse was analyzed irrespective of its location within the brain. Due to missing EKG recordings from 13 participants, optical analyses were limited to a sample size of 112. Optical pulse data were acquired from most of the scalp surface but with varying degrees of density (indexed by the number of channels) depending on the specific study. Study 1 utilized two optical recording montages, each containing 768 channels, for a total of 1536 channels and acquired with one block of recording. Study 2 was acquired with one montage containing 768 channels and acquired with two blocks of recordings. The difference in number of montages used in the two studies did not introduce significant changes in the optical pulse parameters used in this study (see below).

#### 2.2.4 Systemic Pulse Pressure

Participants blood pressure was taken from the brachial artery with an automatic sphygmomanometer placed on the upper arm, at 3 time points during data collection and then averaged together to get more stable estimates of resting diastolic and systolic pressure. Pulse pressure was computed by subtracting diastolic pressure from systolic pressure and served as a systemic measure of cerebrovascular health that is methodologically independent from the cerebral Pulse-DOT measure, thus allowing for external validation of the cerebrovascular measures.

### 2.3 Data Processing

2.3.1 *Orthogonalization of Age and Fitness*

To study the independent contributions of aging and fitness to white matter microstructural integrity, these two variables were orthogonalized. This procedure included three steps. First, the sample was split into three ≈15-year age groups: young (aged 25-40, *n* = 26), middle (aged 41-55, *n* = 33), older adults (aged 56-71, *n* = 68). Within each age group, participants were median split based on their eCRF_SexAdjusted scores into “high fit” and “low fit” categories for their age. Then the groups were recombined for further analysis. The high-fit and low-fit groups did not differ in age, sex, or education level (**Table 1**), but did differ in eCRF_SexAdjusted and its components (Heart Rate, BMI), as planned, and in fluid intelligence.

**Table 1.** Sample characteristics. Means and standard deviations are provided as well as results from 2-tailed t-tests between the low fit and high fit individuals on every sample characteristic.

|  | All (n = 125) |  | Low Fit (n = 61) |  | High Fit (n = 64) |  | t-test 2-tailed |  |
| --- | --- | --- | --- | --- | --- | --- | --- | --- |
| Females: Males | 80:45 |  | 42:20 |  | 39:25 |  |  |  |
|  | Mean | SD | Mean | SD | Mean | SD | *p* |  |
| Age (years) | 53.35 | 13.52 | 54.56 | 13.75 | 52.17 | 13.29 | 0.32 | ns |
| Sex | 0.36 | 0.48 | 0.33 | 0.47 | 0.39 | 0.49 | 0.47 | ns |
| eCRF_SexAdjusted | 9.34 | 2.20 | 7.91 | 1.79 | 10.70 | 1.63 | 1.59E-14 | ** |
| BMI (kg/m<sup>2</sup>) | 27.01 | 5.16 | 29.22 | 5.75 | 24.91 | 3.41 | 2.44E-06 | * |
| Heart Rate (bpm) | 71.19 | 12.73 | 76.89 | 11.54 | 65.75 | 11.42 | 1.22E-06 | * |
| Education (years, 20 max) | 17.15 | 2.39 | 16.90 | 2.58 | 17.39 | 2.19 | 0.34 | ns |
| Fluid Intelligence (n=123) | 105.93 | 10.96 | 103.47 | 12.50 | 108.27 | 8.72 | 0.02 | * |

#### 2.3.2 MRI Preprocessing and Reconstruction

All MRI data were first converted into BIDS format (Gorgolewski et al., 2016). Diffusion weighted images were then run through *QSIPrep* 0.16.0RC3 (Cieslak et al., 2021) for preprocessing and reconstruction, which is based on *Nipype* 1.8.1 (Gorgolewski et al., 2011 Gorgolewski et al., 2018; RRID:SCR_002502), with the parameters listed in the following sections. Much of the text in the following two sections was provided by QSIPrep under a CC0 license so it may be included in a manuscript for the sake of transparency and reproducibility. We made minor changes for clarity.

##### 2.3.2.1 Anatomical data preprocessing

For each participant, the T1-weighted (T1w) image was corrected for intensity non-uniformity (INU) using N4BiasFieldCorrection (Tustison et al. 2010, *ANTs* 2.3.1), and used as T1w-reference throughout the workflow. The T1w-reference was then skull-stripped using antsBrainExtraction.sh (*ANTs* 2.3.1), with OASIS as target template. Spatial normalization to the ICBM 152 Nonlinear Asymmetrical template version 2009c (Fonov et al., 2009, RRID:SCR_008796) was performed through nonlinear registration with antsRegistration (ANTs 2.3.1, RRID:SCR_004757, Avants et al., 2008), using brain-extracted versions of both T1w volume and template. Brain tissue segmentation of cerebrospinal fluid (CSF), white-matter (WM) and gray-matter (GM) was performed on the brain-extracted T1w using FAST (FSL 6.0.5.1:57b01774, RRID:SCR_002823, Zhang, Brady, and Smith, 2001). T1-weighted MPRAGE scans were cropped to include only the head, to resolve errors in brain extraction and registration for three participants (using robustfov in *FSLeyes*; McCarthy, 2023).

##### 2.3.2.2 Diffusion data preprocessing

Any images with a b-value less than 100 s/mm∧2 were treated as a b=0 image. MP-PCA denoising as implemented in *MRtrix3’s* dwidenoise (Veraart et al., 2016) was applied with a 5-voxel window. After MP-PCA, B1 field inhomogeneity was corrected using dwibiascorrect from *MRtrix3* with the N4 algorithm (Tustison et al., 2010). After B1 bias correction, the mean intensity of the DWI series was adjusted so all the mean intensity of the b=0 images matched across each separate DWI scanning sequence.

*FSL’s* (version 6.0.5.1:57b01774) eddy procedure was used for head motion correction and Eddy current correction (Andersson and Sotiropoulos, 2016). Eddy was configured with a q-space smoothing factor of 10, a total of 5 iterations, and 1000 voxels used to estimate hyperparameters. Linear and quadratic models were used to characterize Eddy current-related spatial distortion. q-space coordinates were forcefully assigned to shells. Field offset was separated from subject movement. Shells were aligned post-eddy. Eddy’s outlier replacement procedure was also used (Andersson et al., 2016). Data were grouped by slice, only including values from slices determined to contain at least 250 intracerebral voxels. Groups deviating by more than 4 standard deviations from the prediction had their data replaced with imputed values. Additional data was collected with reversed phase-encode blips, resulting in pairs of images with distortions going in opposite directions. Here, b=0 reference images with reversed phase encoding directions were used along with an equal number of b=0 images extracted from the DWI scans. From these pairs the susceptibility-induced off-resonance field was estimated using a method similar to that described in Andersson, Skare, and Ashburner, 2003. The fieldmaps were ultimately incorporated into the Eddy current and head motion correction interpolation. Final interpolation was performed using the jac method.

Several confounding time-series were calculated based on the preprocessed DWI: framewise displacement (FD) using the implementation in *Nipype* (following the definitions by Power et al., 2014). The head-motion estimates calculated in the correction step were also placed within the corresponding confounds file. Slicewise cross correlation was also calculated. The DWI time-series were resampled to ACPC, generating a preprocessed DWI run in ACPC space with 2.5mm isotropic voxels. *QSIprep* creates HTML summary output files for each participant. These were visually inspected for errors, e.g., inaccurate T1 to MNI registration, and if errors were found, the *QSIprep* pipeline was re-run.^3^

##### 2.3.2.3 Diffusion Reconstruction and Tensor Fitting

Two reconstruction workflows were performed using *QSIprep* 0.16.0RC3, which is based on *Nipype* 1.8.1 (Gorgolewski et al., 2011; Gorgolewski et al., 2018; RRID:SCR_002502). First, *QSIPrep*-preprocessed T1w images and brain masks were used in the reorient_fslstd workflow. This workflow reorients the QSIPrep preprocessed DWI and bval/bvec to the standard *FSL* orientation. Because we were using *FSL* for subsequent analysis, we implemented this workflow. More information can be found here, https://qsiprep.readthedocs.io/en/latest/reconstruction.html#reorient-fslstd

Then, the *QSIPrep*-preprocessed T1w images and brain masks were used in the dsi_studio_gqi workflow (DSI Studio Reconstruction). This implementation reconstructs diffusion orientation distribution functions (ODFs) using generalized q-sampling imaging (GQI, Yeh, Wedeen, and Tseng, 2010) with a ratio of mean diffusion distance of 1.250000. Diffusion tensors (FA, RD, MD, and AD) are also calculated in this workflow and are the main focus of this manuscript. Additional outputs from this workflow (ODFs and streamlines) will be used in future investigations of this dataset. Critically, head motion was also calculated in this pipeline and specifically, framewise displacement was used to assess head motion in the current study (Hoinkiss & Porter, 2022; Kreilkamp et al., 2016). More information can be found here: https://qsiprep.readthedocs.io/en/latest/reconstruction.html#dsi-studio-gqi

Many internal operations of *QSIPrep* use *Nilearn* 0.9.1 (Abraham et al., 2014, RRID:SCR_001362) and *Dipy* (Garyfallidis et al., 2014). For more details of the pipeline, see the section corresponding to workflows in QSIPrep’s documentation https://qsiprep.readthedocs.io/en/latest/.

##### 2.3.2.4 Extraction of Tensors

Fractional Anisotropy (FA) images from *QSIPrep* reconstruction were consolidated into a single folder. Next, a portion of the tract-based spatial statistics (TBSS, (S. M. Smith et al., 2006) pipeline was run in *FSL* (version 6.0.5.1; Smith et al., 2004) using NeuroDesk (v20221216, (Renton et al., 2022) with the instructions provided in the *FSL* tutorial using the recommended parameters (https://fsl.fmrib.ox.ac.uk/fsl/fslwiki/TBSS/UserGuide).^4^

TBSS was used to identify the mean, group-level FA skeleton, i.e., shared white matter voxels across the sample, but rather than computing voxel-based statistics, we implemented an ROI approach. Using the voxels within the group-level FA skeleton ensured that at the participant level, the number of voxels represented within each ROI was equivalent across participants. This ensured that each ROI could be directly compared person-to-person.

ROI masks from the JHU ICBM-DTI-81 white-matter labels atlas (Hua et al., 2008; Wakana et al., 2007) were created for each of the 48 regions in this atlas using the fslmaths function. This atlas is available in *FSL* (https://fsl.fmrib.ox.ac.uk/fsl/fslwiki/Atlases) and has 48 WM tract labels for diffusion MRI tensor maps. These were then applied to the 4D tensor data matrix (N-subjects by X by Y by Z) using custom Matlab 2022a scripts to extract mean FA estimates in each ROI for every participant (The MathWorks Inc., Natick, MA USA). Because many regions are bilateral, but we had no specific hypotheses about laterality, we combined bilateral ROIs into one mean FA estimate (i.e., the superior corona radiata on the left and right were combined into a single “superior corona radiata” mean value). This reduced the number of ROIs to be analyzed to 27.

#### 2.3.3 Optical Measures of Cerebral Arterial Elasticity

Cerebral arterial elasticity was quantified using two pulse-DOT metrics, pulse transit time (PTT) and Pulse Relaxation Function (PReFx, formerly ‘arterial compliance’; Fabiani et al., 2014). These metrics were chosen because they are indices of cerebral arterial elasticity and were found to be related to age and fitness in previous reports. Differences in optical data acquisition between the two studies (i.e., different number of channels in each montage) did not significantly impact the data, so data were combined for all analyses^5^.

PReFx is a measure quantifying pulse shape, which is known to change as a function of the temporal overlap between the forward and backward pressure waves generated during each cardiac cycle. A greater overlap between these two waves occurs due to arterial stiffening, which increases PWV in the segment between the measurement point in elastic arteries, and the reflection point located in small muscular arteries and arterioles (see Izzo & Shykoff, 2001). As such, lower PReFx values index arteriosclerosis, especially in small vessels.

PTT, instead, is the inverse of pulse wave velocity (as distances cannot be easily calculated for cerebral arteries running within the skull) and it is largely a measure of arteriosclerosis in larger elastic arteries. Longer pulse transit times (i.e., slower pulse wave velocities, PVWs) are associated with higher arterial elasticity, whereas shorter pulse transit times (i.e., faster PVWs), indicate reduced elasticity. To calculate PTT, the arterial pulse waveform for each channel was obtained by averaging the AC light intensity time-locked to the peak of the R-wave on the EKG, ensuring that the same pulse cycle was measured at all locations (Fabiani et al., 2014).

To derive PTT and PReFx, optical AC intensity data (i.e., the average measures of the amount of light produced by a specific source and reaching a specific detector during a multiplexed 1.6 ms interval) were normalized, movement corrected (Chiarelli et al., 2015), and low-pass filtered at 10 Hz using a Butterworth filter. As mentioned, for each participant, the location of sources and detectors used for each channel were digitized and co-registered with the corresponding T1w MPRAGE anatomical image (Chiarelli et al., 2015). Only source-detector distances of 2 to 5.5 cm were used in the analyses, as this is a distance range within the adult cortex that provides a sufficient amount of light for reliable calculations, while minimizing the effects of phenomena too superficial to be within the cortex. These data allowed us to apply the 3D reconstruction procedure described in Chiarelli et al. (2015) which makes it possible to image phenomena occurring approximately between 1 and 3 cm in depth within the brain, thereby providing coverage of most of the cerebral cortex. Specifically, an average pulse waveform could be computed for each voxel (and participant) in the head up to this depth, and pulse parameters could be computed for each voxel. Here we employed global measures of cerebral arterial elasticity, obtained by averaging the pulse-DOT parameters across all voxels. Computation of these parameters and averaging across the head was done using in-house software Opt-3d (Gratton, 2000). Note that the odd-even pulse reliability of the optical parameters obtained in this way is very high (*r* > 0.95). In this paper we focus on PTT (reflecting PWV), which is expected to strongly correspond to systolic blood pressure.

### 2.4 Statistical Procedure

Statistical analyses were performed in R version 4.2.1 (www.r-project.org/, R Core Team, 2012) and RStudio (version: RStudio-2022.07.1-554.) A spreadsheet with demographic variables, fitness estimates (eCRF), PTT, and FA mean values for each ROI was created. Then this was imported into R and Spearman’s rho correlations between age, fitness (orthogonal to age), and the age-by-fitness interaction were calculated by mean-centering age and fitness and then multiplying the mean-centered measurements together. Correlations between each ROI and the non-brain variables can be found in **Supp. Table 2**. Spearman’s rho was used to minimize the impact of skewed data on the results.

#### 2.4.1 Principal Components Analysis

To reduce the number of comparisons between FA regions and our dependent variables of interest, we conducted a principal components analysis (PCA) on the ROI FA values. PCA is a data reduction method that transforms high-dimensional data into lower-dimensional data while retaining most of the variability in the original data. It results in a set of principal components (PCs) that are linear transformations of the original data, and these PCs group together variables with shared variance. PCA was computed using the R packages *FactoMineR* (Lê, Josse, & Husson, 2008), *missMDA* (Josse & Husson, 2016), and *pracma* (Borchers, 2023; https://CRAN.R-project.org/package=pracma). The resulting loadings were rotated with a varimax rotation to maximize spread while retaining orthogonality between components. The top three factors were retained for further analysis (henceforth referred to as Midline FA Factor 1, Inferior-Superior FA Factor 2, and Hindbrain FA Factor 3) because they captured 60.19% percent of the variance, and after assessing the scree plot, there was little additional variance to be accounted for by retaining more factors^6^. The brain regions that load onto each retained factor are included in **Table 2** and the regions with loadings greater than 0.30 can be visualized in **Figure 1**. Rotated factor scores at each region for each participant were calculated and retained for further analysis.

**Figure 1.**
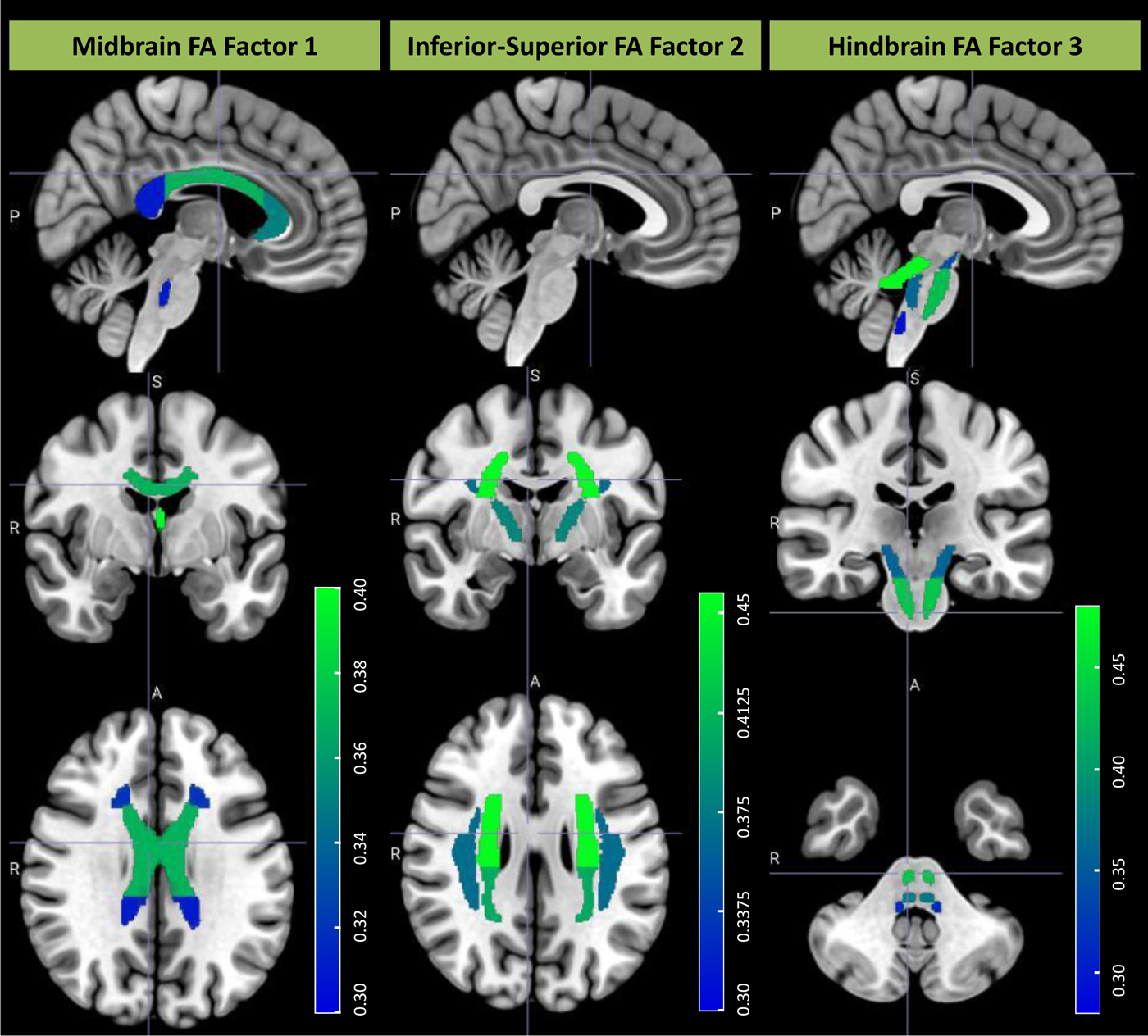
Results of the PCA: Midline FA Factor 1, Inferior-Superior FA Factor 2, and Hindbrain FA Factor 3. Only regions with the highest loadings are visualized (greater than 0.30). The color scale represents the strength of the factor loadings for a particular brain region in the JHU ICBM-DTI-81 white-matter labels Atlas, with the dark blue indicating factor loadings of 0.3 and the light green indicating factor loadings of 0.4 or greater. Hence, the brightest, lightest green regions indicate the white matter region with the largest factor loading for the three PCA factors.

**Table 2.** Rotated Factor Loadings from the PCA.

| Midline FA Factor 1 Loadings |  | Inferior-Superior FA Factor 2 Loadings |  | Hindbrain FA Factor 3 Loadings |  |
| --- | --- | --- | --- | --- | --- |
| Fornix | 0.376 | Superior Corona Radiata | 0.459 | Superior Cerebellar Peduncle | 0.471 |
| Corpus Callosum - Body | 0.354 | Posterior Corona Radiata | 0.394 | Corticospinal Tract | 0.427 |
| Corpus Callosum - Genu | 0.329 | Posterior Limb of Internal Capsule | 0.356 | Medial Lemniscus | 0.359 |
| Posterior Thalamic Radiation | 0.305 | Superior Longitudinal Fasciculus | 0.346 | Middle Cerebellar Peduncle | 0.328 |
| Corpus Callosum - Splenium | 0.301 | Retrolenticular Part of Internal Capsule | 0.306 | Inferior Cerebellar Peduncle | 0.297 |
| Tapetum | 0.300 | Superior Fronto-occipital Fasciculus | 0.247 | Cingulum - Hippocampus | 0.217 |
| Anterior Corona Radiata | 0.271 | External Capsule | 0.206 | Cerebral Peduncle | 0.222 |
| Fornix Cres/Stria Terminalis | 0.232 | Sagittal Stratum | 0.187 | Internal Capsule | 0.201 |
| Cingulum - Cingulate Gyrus | 0.225 | Internal Capsule | 0.179 | Pontine Crossing Tract | 0.192 |
| Sagittal Stratum | 0.198 | Anterior Corona Radiata | 0.133 | Posterior Limb of Internal Capsule | 0.146 |
| External Capsule | 0.152 | Superior Cerebellar Peduncle | -0.118 | Fornix Cres/Stria Terminalis | 0.127 |
| Pontine Crossing Tract | 0.102 | Fornix | -0.211 |  |  |
| Corticospinal Tract | -0.123 |  |  |  |  |
| Superior Cerebellar Peduncle | -0.134 |  |  |  |  |
| Posterior Limb of Internal Capsule | -0.169 |  |  |  |  |

#### 2.4.2 Correlational Analyses

Correlations between the three retained Factors, age, fitness, their interaction, PTT, sex, and head motion (quantified using mean absolute value framewise displacement per person) were computed and their raw correlations are reported in **Supp. Figure S1**. We determined that head motion was significantly correlated with age (*rho*(123) = 0.27, *p* = 0.002), fitness (*rho*(123) = -0.28, *p* = 0.0016 ), Midline FA Factor 1 (*rho*(123) = -0.33, *p* = 0.0002), and marginally correlated with Hindbrain FA Factor 3 (*rho*(123) = -0.16, *p* = 0.08). Sex was also correlated with pulse transit time (greater for females, *rho*(123) = -0.24, *p* = 0.01), Midline FA Factor 1 (greater for females, *rho*(123) = -0.23, *p* = 0.01), and Inferior-Superior FA Factor 2 (greater for females, *rho*(123) = 0.18, *p* = 0.04). Given that our primary aim was to assess the relationship between white matter integrity, age, fitness, and cerebrovascular health across both sexes and free from head motion, correlations between the PCA factors and the biological dependent variables are reported after *partialling out* head motion and sex.

#### 2.4.3 Piecewise Regression

Our primary interest was in how vascular health impacts the relationship between age and white matter microstructure. As mentioned in the introduction, declines in vascular health start becoming evident in middle-age. As such, decline in variables that are likely to depend on the health of the vasculature is expected to accelerate as arteriosclerosis increases. Therefore, we also conducted piecewise regression/bilinear analyses between age and the factors retained from the PCA to determine if any white matter factors showed bilinear trends. In this analysis, two lines joined by a single breakpoint – hypothesized to occur slightly after the typical age of vascular changes (∼55) – were fit to the data. This allows us to estimate the age at which the pre- and post-breakpoint slopes differ from one another. The *segmented* package in R was used to complete this analysis (Muggeo, 2008). We employed the (pseudo) score statistics (Muggeo, 2016) via the ‘pscore.test()’ function to test for a non-zero difference in slope parameter of the bilinear relationship. Simulation studies have shown that this test is more statistically powerful than the alternative Davies test (Davies, 1987) when only one breakpoint is being tested (Muggeo, 2016), as we are computing in the current work.

#### 2.4.4 Mediation Models

In four simple mediation analyses, we investigated the links between aging or fitness and white matter integrity through the mediating role of arterial elasticity, measured both *cerebrally* with Pulse-DOT and *peripherally* with pulse pressure. Using the *PROCESS* macro developed for R (Hayes, 2017), we tested the following simple mediation models. Mediation models were only conducted between the significant relationships found in the correlational analyses:

1. Age → Pulse Transit Time (cerebral measure) → Midline FA in Factor 1
2. Fitness → Pulse Transit Time (cerebral measure) → Hindbrain FA in Factor 3
3. Age → Pulse Pressure (peripheral measure) → Midline FA in Factor 1
4. Fitness → Pulse Pressure (peripheral measure) → Hindbrain FA in Factor 3

Results from validation models with pulse pressure (Models 3 & 4) are included in the supplementary materials (described and specified below).

In line with the correlational analyses, sex and head motion were included as covariates in each of the four mediation models. All models without these covariates can be found in the Supplemental Materials (**Supp. Figures S2-5**). This analytic strategy was chosen for its directional path approach, which is the best option here, given the limitations of a cross-sectional study. We tested the indirect effect by using a bias-corrected bootstrap approach with 5000 samples (Hayes & Scharkow, 2013; Preacher & Kelley, 2011).

#### 2.4.5 Sex Analyses

As a secondary aim, and in line with the National Institute of Health’s initiative (Arnegard et al., 2020; Miller et al., 2017), we assessed the impact of sex on the relationship between age, fitness, and white matter integrity. Please note that we use self-reported sex (i.e., how individuals self-identify, not necessarily their genetic makeup). There were only two self-reported sexes (male, female), as no participants in our study identified as non-binary or did not respond.

All reported correlations are corrected for head motion (partial Spearman correlation). Our sample contained 80 females and 45 males, so the correlations reported below are not balanced by sex. Given the reduced sample size and therefore statistical power, mediation models were not computed for sex subsamples. First, we assessed whether white matter integrity for each Factor score differed by sex. Then, we split the sample into males and females and repeated the above analyses.

## 3. Results

### 3.1 Correlational Analyses

Given the primary interest in assessing the independent effects of age and fitness on white matter integrity, we began by assessing correlations between these variables on the PCA factors we derived. Correlations are summarized in **Table 3**. As a reminder, all correlations with PCA factors are corrected for head motion and sex (and therefore have N-4 degrees of freedom). Correlations were corrected for multiple comparisons within each ‘non-brain’ variable (i.e., Age, Fitness, Interaction) using Holm’s method. Age was significantly correlated with Midline FA Factor 1, *rho*(121) = -0.39, *p* = 0.027, indicating that as individuals get older, FA is reduced in the corpus callosum and fornix, among others. Age was not significantly related to Inferior-Superior FA Factor 2, *rho*(121) = -0.17, *p* = 0.09, and Hindbrain FA Factor 3, *rho*(121) = -0.07, *p* = 0.47. Fitness, which is orthogonal to age, on the other hand, was significantly related to Hindbrain FA Factor 3, *rho*(121) = 0.24, *p* = 0.03, indicating that FA in deep brainstem regions some of which connect the cortex with other regions such as the cerebellum and the spinal cord, increases with increasing CRF levels. Fitness was not significantly related to Midline FA Factor 1, *rho*(121) = 0.14, *p* = 0.14, and Inferior-Superior FA Factor 2, *rho*(121) = 0.16, *p* = 0.14. Lastly, the interaction between aging and fitness was significantly related to Midline FA Factor 1 only prior to multiple comparisons correction, *rho*(120) = 0.18, *p* = 0.04, but not after (*p* = 0.12). Although this effect was marginal, the theoretical implications of this interaction would support an moderating effect of fitness on age-related changes in white matter integrity, leading us to probe it further. As visualized in **Figure 2**, higher fit individuals generally have a slower rate of change in white matter integrity compared to lower fit individuals, suggesting that fitness positively offsets the effect of aging in the white matter areas connecting the hemispheres. Computing the Johnson-Neyman Interval (Bauer et al., 2005; Esarey & Sumner, 2018; P. O. Johnson & Fay, 1950) using the *interactions* package in R (Long, 2022) indicates that after the age of 55.37 years, the effect of fitness on age is significant: lower fit individuals have a steeper slope than higher fit individuals beginning in middle-age and thereafter. Interestingly, age 55 is during the range in which vascular decline beings (pulse pressure increase, Izzo & Shykoff, 2001) indicating that fitness may serve a protective role during this vulnerable period.

**Table 3.** Correlations between FA Factors and non-brain variables. +p < .1, *p < .05, Significance denotation is after correction for multiple comparisons.

|  | Age | Fitness | Interaction |
| --- | --- | --- | --- |
| **Midline FA Factor 1** | -0.39 * | 0.14 | 0.18 |
| **Inferior-Superior FA Factor 2** | -0.17 + | 0.16 | 0.16 |
| **Hindbrain FA Factor 3** | -0.07 | 0.24 * | 0.12 |

**Figure 2.**
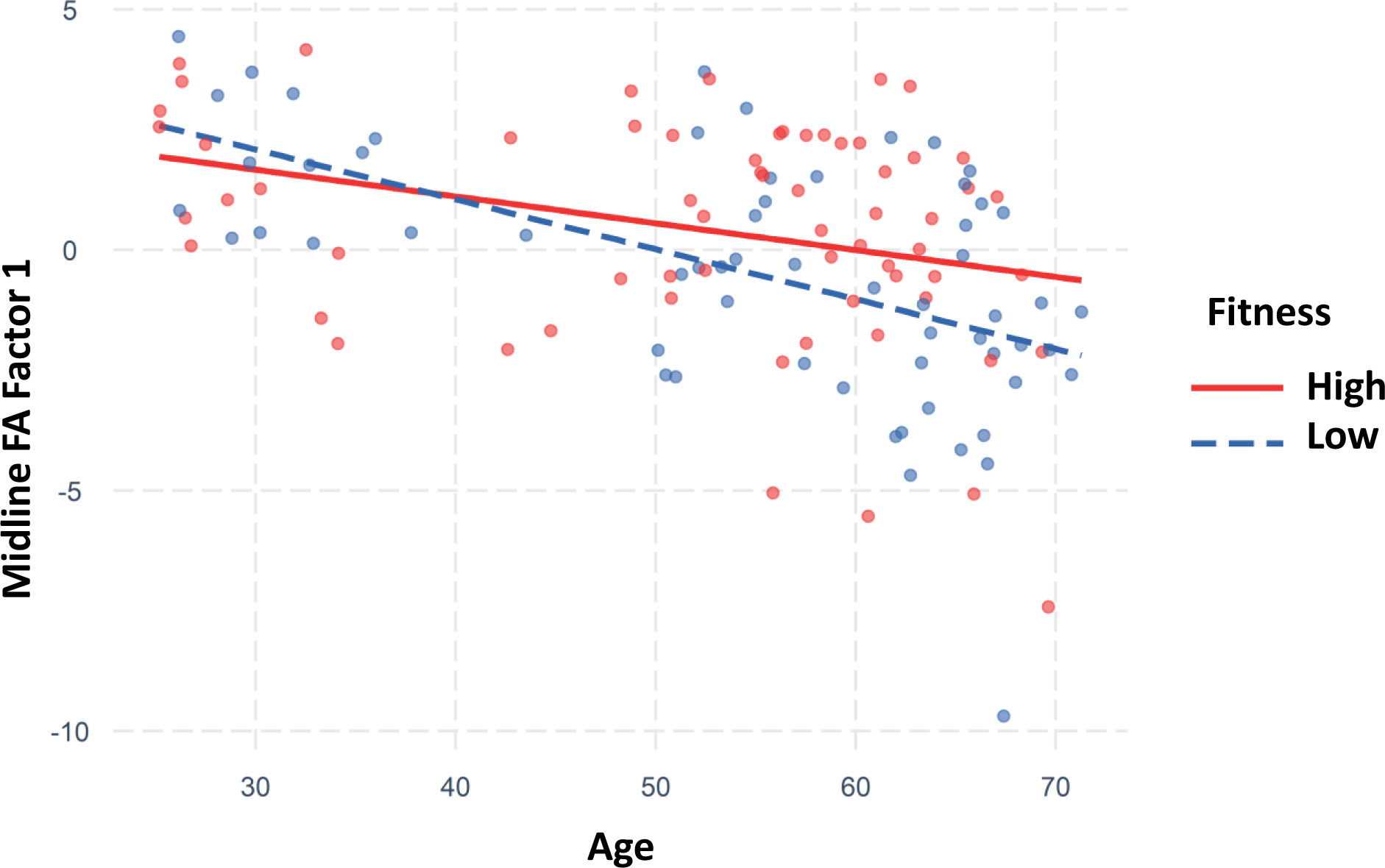
Interaction between age and fitness on Midline FA Factor 1. Red dots indicate individuals with high fitness and their corresponding trendline in red. Blue dots are those with low fitness and their corresponding trendline in dashed-blue. As a reminder, fitness was median split within 3 age ranges: 25-40, 41-55, and 56-72. Generally, individuals with higher fitness show a slower rate of change in their Midline FA captured in Factor 1 compared to those with lower levels of fitness and this effect was statistically reliable after age 55.

### 3.2 Piecewise Regression/Bilinear Analysis

To further understand the relationship between age and Midline FA Factor 1, we computed a bilinear regression analysis in line with the prediction that FA would have non-linear relationships with age, due to the middle-age acceleration in vascular problems (Najjar et al., 2005; Sun, 2015). The model identified a significant breakpoint at age 64.32 (SE = 2.42, p < 0.001, 95% CI [59.54, 69.12]), in which the regression lines before and after this age (β1 = -0.06, 95% CI [-0.09, -0.02], β2 = -0.44, 95% CI [ -0.93, 0.05]) have significantly different slopes from one another (**Figure 3**). However, do note that only the pre-breakpoint estimate β1 was significant, indicating only small changes in FA in these primarily midline regions up until mid-life. The post-breakpoint slope, although clearly steeper, was non-significant, likely due to a smaller number of participants older than age 64. This points to the need for further studies with a larger sample of adults older than 64. Importantly, the onset of accelerated vascular dysfunction is typically earlier than the breakpoint age reported for the FA-age slope, with increases in pulse pressure accelerating between 50-60 years (Izzo & Shykoff, 2001). Using a Granger-like causality logic, this may suggest that the acceleration of FA loss with age could be a delayed consequence of cerebrovascular decline.

**Figure 3.**
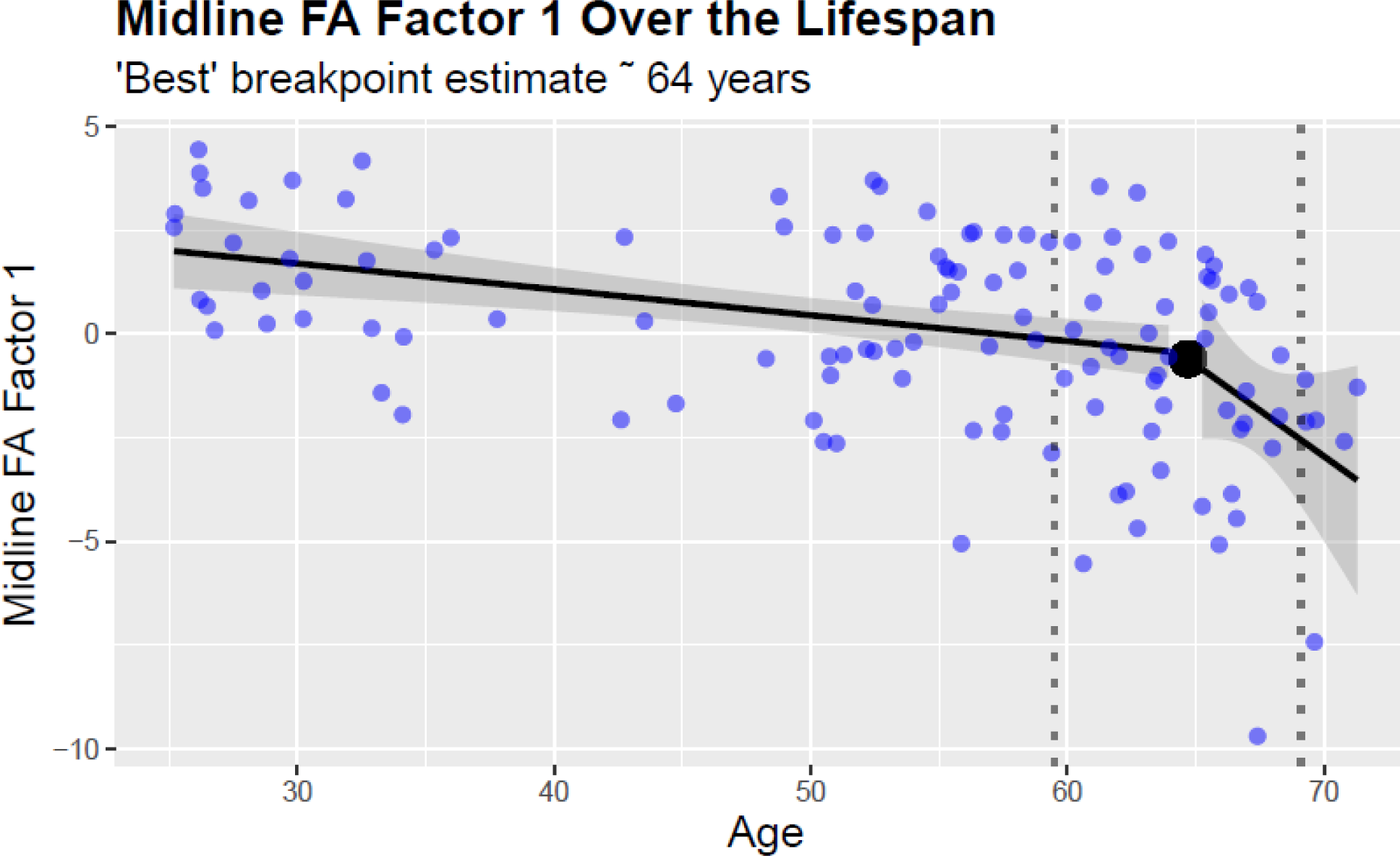
Bilinear analysis of Midline FA Factor 1 across the adult lifespan. The slopes of the 2 regression lines are significantly different from one another before/after age 64, indicated by the large black dot. The dashed lines indicate a 95% bootstrapped confidence interval about the breakpoint of 64.

Age did not have a bilinear relationship with the other two factors from the PCA. Although a breakpoint was identified between age and Inferior-Superior FA Factor 2 at 34.09 years, it was not significant (SE = 4.84, p = 0.35, 95% CI [24.51, 43.68]) and neither were the pre- or post-breakpoint regression lines (β1 = -0.24, 95% CI [-0.57, 0.08], β2 = -0.0005, 95% CI [ -0.05, 0.05]). Similarly, a breakpoint was identified between age and Hindbrain FA Factor 3 at 26.77 years, but it was not significant (SE = 0.69, p = 0.94, 95% CI [25.41, 28.13]) and neither were the pre- or post-breakpoint regression lines (β1 = -2.07, 95% CI [-5.01, 0.87], β2 = 0.004, 95% CI[ -0.031, 0.038]). The figures for these can be found in the Supplemental Materials (**Supp. Figures S6-7**). These models indicate that only the Midbrain FA Factor 1, strongly related to age, show bilinear relationships, and that the bilinear shape is likely related to the aging process itself.

### 3.3 Mediation Models

#### 3.3.1 Age → Arterial Health → Midline FA Factor 1

Correlations between age, fitness, their interaction, sex, PReFx, and the three retained factors, were computed and are reported in **Supp. Table 1**. As can be seen, PReFx was negatively correlated with age and positively correlated with fitness, as reported by previous studies. However, no significant correlations with the PCA factors were apparent. Therefore, only PTT was used as an estimate of cerebral arterial stiffness for the mediation models.

To complement the prior analyses, we conducted several simple mediation path analyses to test the hypothesized relationships between age or fitness, arterial health (cerebrally derived pulse transit time, or peripherally derived pulse pressure), and white matter integrity (Midline and Hindbrain FA in Factors 1 and 3). As a reminder, 112 participants from the full sample had complete optical data. Mediation models were only conducted between the significant relationships found in the correlational analyses. First, we tested the indirect effect of age on Midline FA Factor 1 via Pulse Transit Time (n = 110, **Figure 4**). Based on a bias-corrected bootstrapped 95% confidence interval, the indirect effect (a x b) was significant, suggesting that age-related declines in fractional anisotropy in regions such as the corpus callosum and fornix are partially explained by cerebral arterial health. This indicates that, although age is associated with declines in both FA and cerebral arterial health, having higher PTT (indicative of healthy/elastic arteries and a slower pulse wave velocity) may be at least partially protective against reductions in midline FA across the lifespan.

Briefly, when systemic pulse pressure is the mediator in the above model instead of the cerebral PTT, the indirect effect (a x b) is not significant (see **Supp. Figure S8**). Despite the averaging of three blood pressure measurements, automatic digital sphygmomanometers are susceptible to measurement error (Shahbabu et al., 2016). In addition, we assume that pulse pressure measured from the brachial artery will be less sensitive to cerebral arterial changes, potentially accounting for the lack of mediation. It should be noted, however, that if the same analysis was repeated without including head motion and sex as covariates, pulse pressure would mediate the relationship as in Model 1, **Supp. Figure S4**).

**Figure 4:**
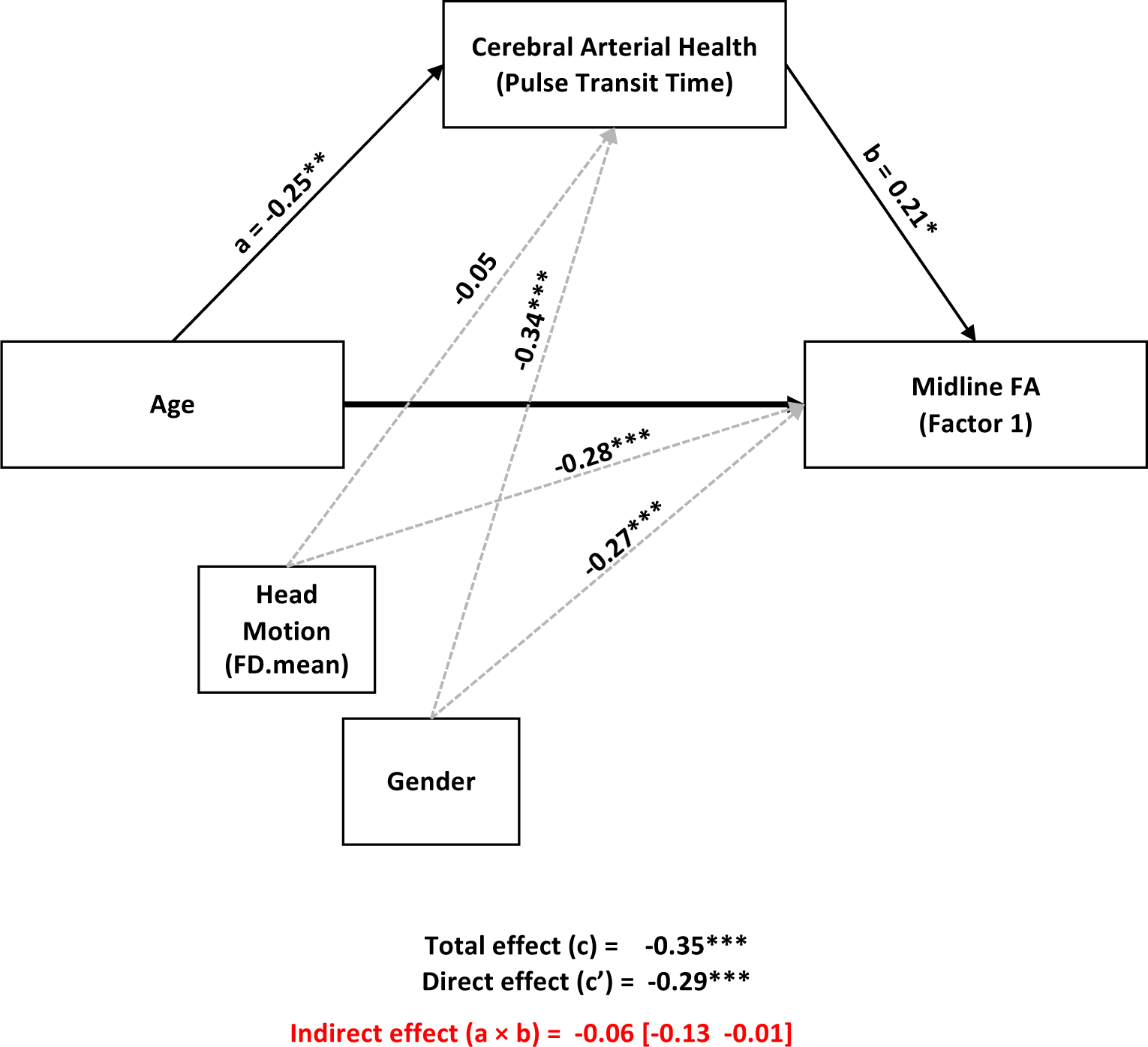
Mediation model of Age on Midline FA Factor 1 via Cerebral Arterial Health measured with Pulse Transit Time. Head motion and Sex are covariates, as indicated by the dashed gray lines. The indirect effect is significant, as indicated by a bootstrapped 95% confidence interval that does not include zero. Standardized coefficients are reported. *p > .05, **p > .01, ***p > .001

#### 3.3.2 Fitness → Arterial Health → Hindbrain FA Factor 3

Next, we tested the indirect effect of fitness on Hindbrain FA Factor 3 via Pulse Transit Time (n = 110, **Figure 5**). Based on a bias-corrected bootstrapped 95% confidence interval, the indirect effect (a x b) was significant, suggesting that the relationship between improved fitness and preserved white matter integrity in brainstem regions is partially due to improved cerebral arterial health. Higher fitness levels are associated with greater PTTs (i.e., healthier/elastic arteries), which in turn are related to preserved white matter integrity, supporting the mediating role of the vasculature in the preservation of white matter. This relationship was evident irrespective of participant age.

**Figure 5.**
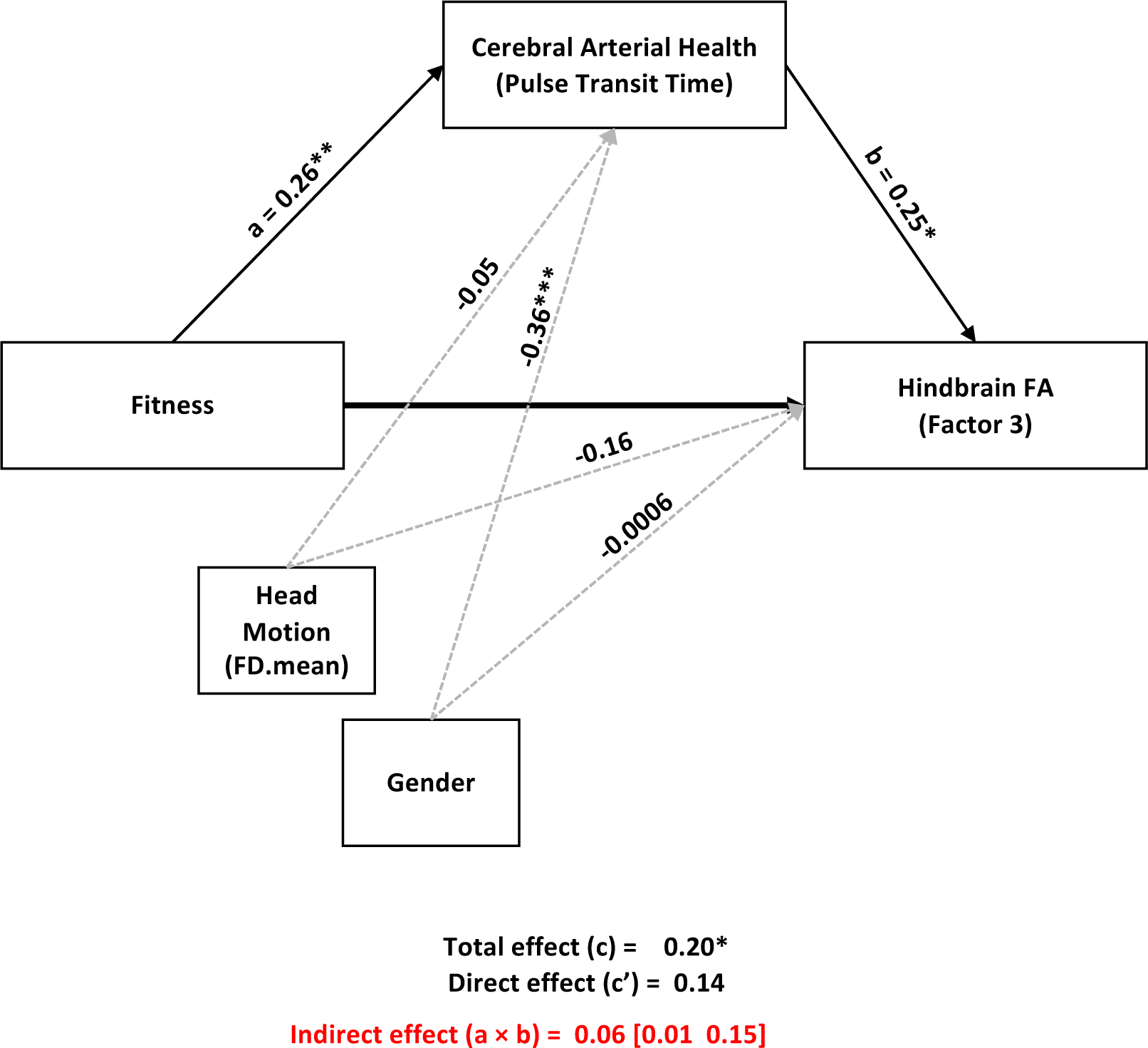
Mediation model of Fitness on Hindbrain FA Factor 3 via Cerebral Arterial Health measured with Pulse Transit Time. Head motion and Sex are covariates, as indicated by the dashed gray lines. The indirect effect is significant. Standardized coefficients are reported. *p > .05, **p > .01, ***p > .001

Briefly, when systemic pulse pressure was used as a mediator in the above model, the indirect effect (a x b) was not significant **(Supp. Figure S9-10)**. Again, although sex accounted for a large amount of variance in pulse pressure, pulse pressure was not a significant mediator prior to including sex and head motion as covariates (**Supp. Figure S5**). This suggests that although the deep brainstem tracts are positively influenced by higher levels of CRF, systemic measures of arterial health do not account for substantial variance in this relationship. It may be that systemic measures of arterial health are not sensitive to such deep white matter tracts, whereas cerebral measures (Pulse-DOT) are.

### 3.4 Sex Analyses

As a reminder, all correlations reported are corrected for head motion (partial Spearman correlation with N-3 degrees of freedom). Given the reduced sample size and therefore statistical power, mediation models were not computed for the sex subsamples.

#### 3.4.1 Relationship between sex and FA Factors

Firstly, we assessed the relationships between sex and the FA PCA Factors. Sex was significantly related to Midline FA Factor 1 such that there was greater FA in females in the corpus callosum/fornix, primarily, compared to men, *rho*(122) = -0.27, *p* > 0.01. Sex was also significantly related to Inferior-Superior FA Factor 2, such that there was greater FA in males in regions such as the superior and inferior corona radiata compared to females, *rho*(122) = 0.17, *p* = 0.05. Sex and Hindbrain FA Factor 3 were not related meaningfully, *rho*(122) = -0.08, *p* = 0.38, indicating that FA in the deep brainstem tracts does not vary by sex.

#### 3.4.2 Age, Fitness, and Their Interaction with Sex

Next, we assessed the correlations reported for the full sample in section 3.1 in females and males separately (**Table 4**). Although fitness was orthogonalized based on participant age, fitness was well balanced in females (low fit = 41, high fit = 39) and fairly well balanced in males (low fit = 20, high fit = 25). In line with the full sample correlations, age was significantly related to Midline FA Factor 1 in females, *rho*(77) = -0.35, *p* < 0.05, and in men, *rho*(42) = -0.45, *p* < 0.01. Prior to correction for multiple comparisons fitness was positively associated with Hindbrain FA Factor 3 in females [*rho*(77) = 0.23, *p* = 0.04], but not after, (*p* = 0.12), and marginally related in males before correction [*rho*(42) = -0.29, *p* = 0.06], but not after (*p* = 0.18). Given the smaller sample size in men, this is in line with the full sample correlations. The interaction between age and fitness was not significantly correlated with any FA Factor in females or men. No other correlations were significant in the sex subsamples. Taken together, these results indicate that the pattern of effects between age and white matter integrity across the brain impacts females and males similarly and that the fitness effects are not as strong as the age effects.

**Table 4.**
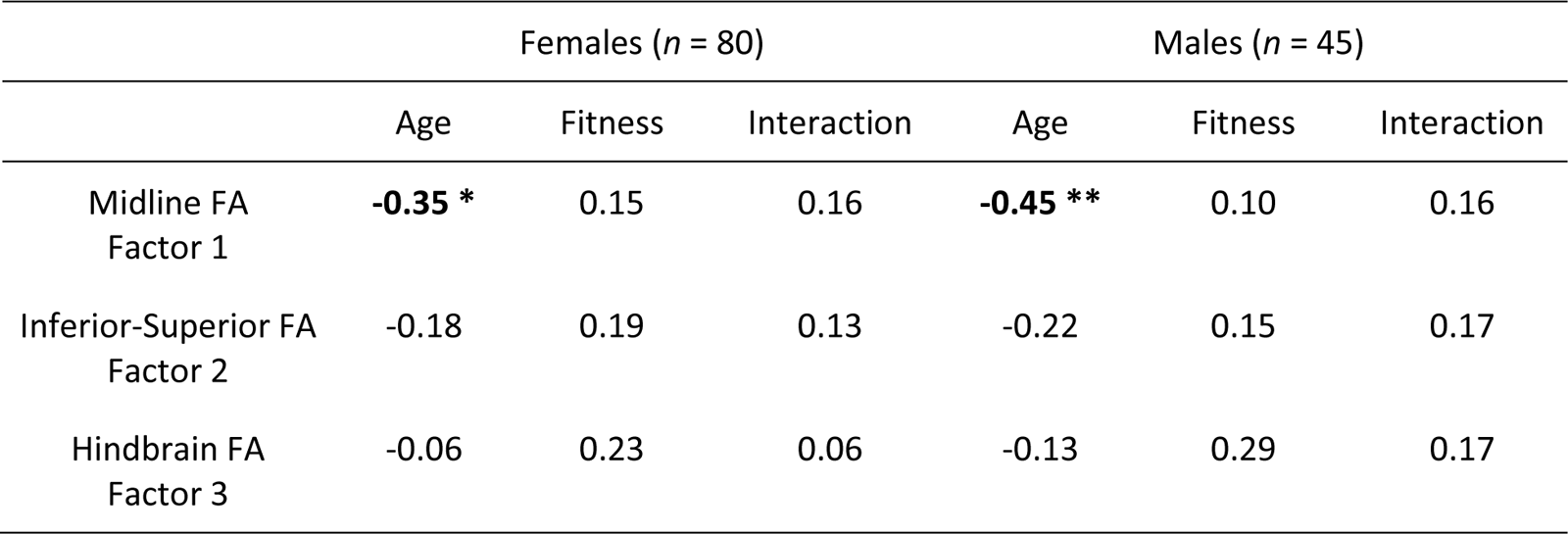
Correlations of Factors in Females and Males. p-values are corrected for multiple comparisons using Holm’s method, *p < .05, **p < .01.

#### 3.4.3 Exploratory Sex Analyses

Given that the magnitude of the correlation between age and Midline FA Factor 1 in males was greater than the effect in females, we sought to understand whether there was an interaction between age and sex for this variable (**Figure 6**). The interaction itself was marginally significant (*p* = 0.065); computing the Johnson-Neyman Interval (Johnson & Fay, 1950) indicates that above the age of 42.21 years, the effect of sex on age is significant: males have a steeper slope than females after this age.

**Figure 6.**
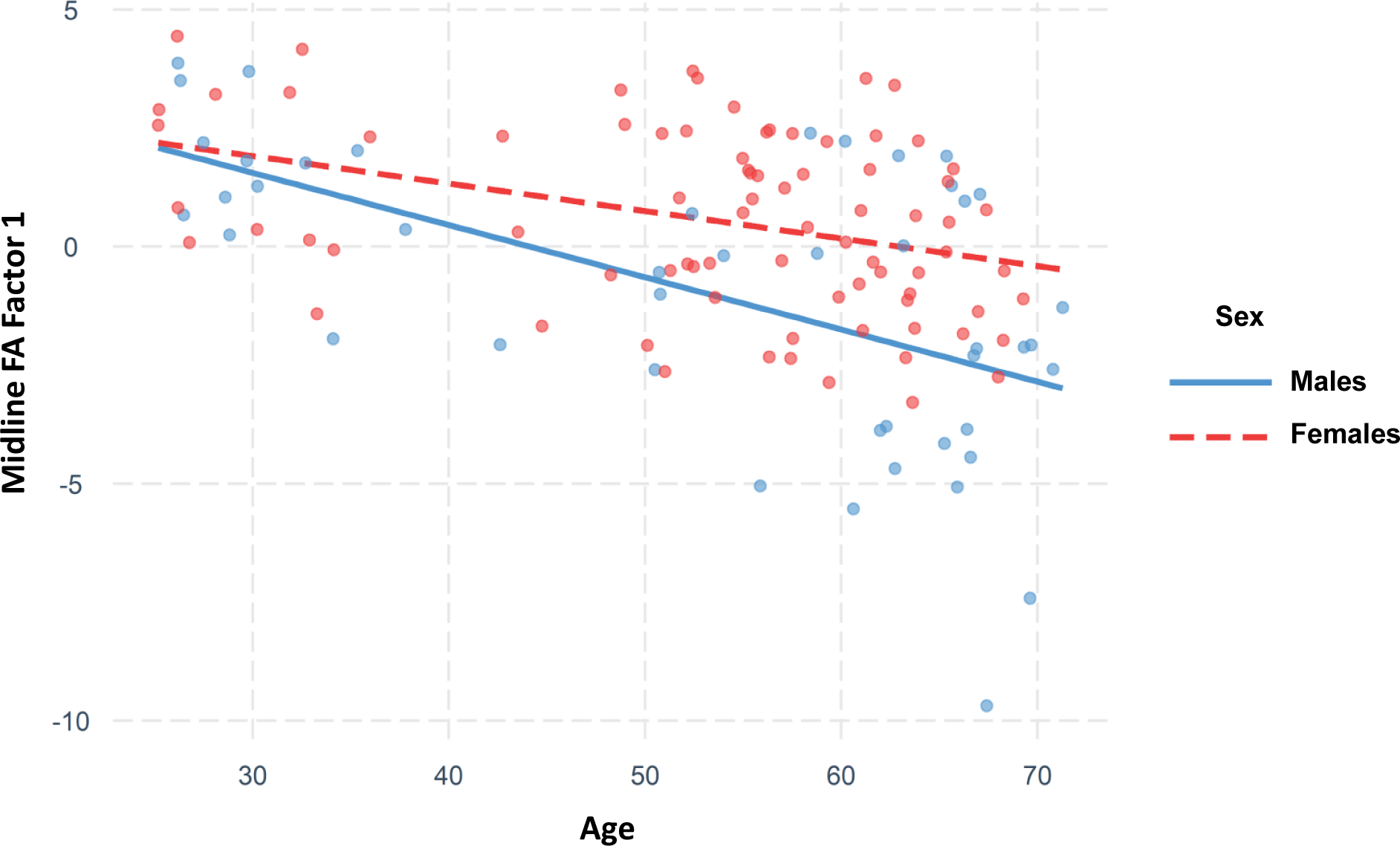
Marginally significant interaction between Age and Sex on Midline FA Factor 1. Red dots indicate females and their corresponding dashed red trendline. Blue dots indicate males and their corresponding solid blue trendline. Generally, females show a slower rate of change in their Midline FA captured in Factor 1 compared to men, across the lifespan.

## 4. Discussion

In this work, we have demonstrated that age and fitness differentially impact white matter integrity across the brain in a cross-sectional sample covering a large portion of the adult lifespan, and that cerebrovascular health acts as a mediator in these relationships. Fractional anisotropy in the fornix and corpus callosum, as well as other regions such as the cingulum of the cingulate gyrus (Midline FA Factor 1), was correlated with age in the predicted direction, with decreased integrity with increasing age. However, this relationship was bilinear in nature, with accelerated declines after age 64. Fractional anisotropy in deep brainstem regions, including the cerebellar peduncle, corticospinal tract, and medial lemniscus (Hindbrain FA Factor 3), was uniquely correlated with fitness, such that higher fit individuals, irrespective of their age, had more preserved white matter integrity in these regions. These two effects were similar in females and males, although age-related white matter decline was larger in males. Perhaps most interestingly, age and fitness had an interactive effect on regions such as the fornix and corpus callosum (Midline FA Factor 1), such that age-related reductions in white matter integrity were less pronounced in higher fit compared to lower fit individuals. Cerebrovascular health partially mediated the relationship between age and white matter integrity, and also between fitness and white matter integrity, highlighting the importance of vascular phenomena on neuroanatomical outcomes.

Our primary aim concerned identifying which regions of the white matter, if any, were independently related to aging compared to fitness. By orthogonalizing age and fitness in conjunction with the use of principal component analysis to group white matter ROIs into factors, we found that age was strongly related to FA in the fornix and corpus callosum (Midline FA Factor 1), but this age effect interacted with fitness. These data do not support a unique effect of age on these midline white matter regions, but rather that these regions are sensitive to fitness in older adults. Indeed, after age 55, fitness appears to influence white matter integrity to a greater extent than before, with a smaller effect of age, i.e., smaller loss of FA in higher fit older adults. These results are in line with research indicating that a) white matter integrity in the corpus callosum declines with age (i.e., Lebel et al., 2012; Ota et al., 2006; Pfefferbaum et al., 2006), and also b) that the integrity of the corpus callosum is positively correlated with fitness and physical activity (S. M. Hayes et al., 2015; N. F. Johnson et al., 2012; Kim et al., 2020; Liu et al., 2012; Mendez Colmenares et al., 2021; Oberlin et al., 2016; Opel et al., 2019; Strömmer et al., 2020; Tarumi et al., 2021a), as is the fornix (Burzynska et al., 2017; Oberlin et al., 2016; J. C. Smith et al., 2016; Tarumi et al., 2021b). The results extend previous work by showing that fitness does not universally impact white matter integrity across the lifespan; instead, the beneficial effects of fitness emerge in middle age (see Johnson et al., 2020, for convergent findings when measuring white matter hyperintensity volume). It should be noted that fitness alone was not related to white matter integrity in the fornix and corpus callosum, it was only related in conjunction with aging. This is in line with previous findings from longitudinal work indicating that older individuals who undergo a fitness regimen generally see improved FA in these regions (Burzynska et al., 2017; Mendez Colmenares et al., 2021; Voss et al., 2013 but see Clark et al., 2019 for null effects), but that fitness interventions in younger adults do not yield such FA improvements (Lehmann et al., 2020). It may also explain the null findings from Predovan et al. (2021) in which a fitness intervention did not yield significant improvements in FA in adults across the lifespan.

Given that fitness seems to impact white matter integrity differentially across adulthood, analyzing all ages together or all regions (global FA) may not detect fitness effects.

It is important to note that fitness was *uniquely* related to white matter integrity in the brainstem and cerebellum, particularly the cerebellar peduncle, corticospinal tract, and medial lemniscus (Hindbrain FA Factor 3), irrespective of participant’s age. These findings support a small collection of studies reporting similar results in very large samples (> 800) of 45-year-olds (d’Arbeloff et al., 2021) and younger adults (Callow et al., 2022; Opel et al., 2019). Given that these white matter regions connect the cerebrum, brainstem, cerebellum, and spinal cord to coordinate bodily movements, it is not surprising that individuals who are higher fit may spend more time coordinating and planning movements, regardless of their age. At face value, this finding seems to support a unique effect of fitness – having improved fitness presumably because of greater motor activity relates to improved white matter integrity in tracts involved in this activity (consistent with a use-it-or-lose-it logic and the animal studies on experience-dependent information storage, i.e., Greenough et al., 1987; Grossman et al., 2003).

According to the cascade model of neurocognitive aging (Kong et al., 2019; Villeneuve & Jagust, 2015), changes in the (cerebro)vascular system may have direct impact on brain structure, particularly beginning in middle age. In this study, we leveraged pulse-DOT technology to quantify cerebrovascular health and used this metric, along with peripherally derived pulse pressure, as mediators in the aforementioned relationships. Arterial health measured cerebrally with PTT mediated the relationship between age and white matter integrity in the fornix and corpus callosum (Midline FA Factor 1). Critically, this association had not been tested with *cerebral* measures, although Hoagey et al. (2021) identified blood pressure as a mediator between age, white matter integrity and cognition and Fuhrmann et al., (2019) found that both blood pressure and age contributed to white matter integrity in most white matter tracts using complex structural equation models. We found that as age increases, arterial health declines, resulting in decreased FA, which is in line with Hoagey et al.’s (2021) model using peripheral arterial measurements. The health of the white matter in these regions was related to age *via* the cerebral vasculature, suggesting that age-related declines in the corpus callosum and fornix (Midline FA Factor 1) may have vascular antecedents.

Additionally, cerebral arterial health as indexed by PTT mediated the relationship between increased fitness and improved white matter integrity in the brainstem regions, indicating that cerebrovascular health accounts for a substantial proportion of the variance in the fitness and FA relationship. This effect was evident irrespective of participant’s age, indicating that having higher levels of fitness benefits the brainstem white matter across the adult lifespan. It also strongly supports an indirect effect of fitness, whereby engaging in physical activity results in healthier arteries, thereby preserving the brainstem white matter. To our knowledge, this effect has not been previously reported.

Together, these mediation models suggest a critical role of cerebrovascular health in brain aging and fitness effects on white matter integrity. Although age exerts a negative effect on white matter integrity and fitness exerts a positive effect, cerebrovascular health partially explains both effects. This is in line with previous evidence indicating that cerebrovascular health mediates the relationship between age and white matter signal abnormalities (Tan et al., 2019) and that fitness mediates the relationship between age and cerebral blood flow (Zimmerman et al., 2014)—the latter implicitly suggesting a vascular component. Similar models are in line with the cascade model of neurocognitive aging indicating that white matter integrity mediates the relationship between fitness (Oberlin et al., 2016), systolic blood pressure (Acosta et al., 2023), or arterial pulsatility (Conley et al., 2020) and cognitive performance (see also Bowie et al., submitted). In these, the vasculature impacts tissue health, similarly to the current findings, which then has downstream impacts on cognition.

As noted, we unexpectedly found no significant relationships between PReFx and the FA factors. At first glance this would suggest that arteriosclerosis in large arteries may impact FA more than cerebral small vessel disease (see Bowie et al., submitted). However, such inference would be based on accepting the null hypothesis, for which may not have sufficient power. Additional work is needed to further explore the possible dissociation between PReFx and PTT measures of arterial status.

A secondary set of analyses explored how these relationships varied based on self-reported sex. The correlational findings are replicated in each sex, indicating that males and females have roughly similar trajectories between age, fitness, and white matter integrity. Males tend to fare slightly worse across the lifespan in regions such as the fornix and corpus callosum (Midline FA Factor 1). This may be due to the vasoprotective effects of estrogen, specifically estradiol that females have up until menopause (Gilligan et al., 1994; Hurn et al., 1995; Rossi et al., 2011). We did not have sufficient power to test the mediation models separately in these groups; this should be explored in future studies.

Two methodological choices made in the current study should be noted. First, the use of principal components analysis in conjunction with white matter FA ROIs has not been utilized extensively (but see (Fletcher et al., 2016) for a complementary gray matter study and (M. A. Johnson et al., 2014) for a related approach). PCA is a useful tool to reduce the dimensionality of the data as well as grouping similar regions together. Second, we quantified framewise displacement, a measure of head motion in the scanner (Power, 2014; Power et al., 2012) and assessed its relation to our variables of interest. In fMRI studies, age is known to be strongly correlated with head motion (Geerligs et al., 2017; Saccà et al., 2021) and samples including older adults require stricter criteria when doing resting-state functional connectivity analysis (Kong et al., 2019; C. Gratton et al., 2020). However, the effect of head motion on dwMRI metrics has been studied less extensively (but see, (Hoinkiss & Porter, 2022; Kreilkamp et al., 2016), and may be a potential confound. In our study, although head motion was correlated with age and some of the PCA factors, our correlations of interest remained reliable even after correcting for head-motion. Future work in aging samples should take care to quantify head motion and assess its impact on dwMRI outcomes.

### 4.1 Limitations and Future Directions

To extend this work, several improvements could be implemented and new avenues explored. While we report here on the widely used diffusion tensor metrics, more sophisticated modeling procedures now yield complementary metrics that aim to better reflect specific aspects of biophysical reality in white matter (i.e., metrics from model-free generalized q-space imaging using orientation distribution functions (Yeh et al., 2010)); multi-shell multi-tissue constrained spherical deconvolution (Jeurissen et al., 2014); Neurite Orientation Dispersion Density Imaging (NODDI, Kamiya et al., 2020; Zhang et al., 2012)). These newer methods use b-value-specific information about diffusion signal in different tissues (Jeurissen et al., 2014) and attempt to explicitly model diffusion in different compartments – intracellular (axons & dendrites), glia (cell bodies, glial cell, ECM, vasculature), and extracellular isotropic diffusion (free water). With sufficiently sophisticated diffusion weighted imaging acquisitions, these methods provide information that can tease apart changes seen in FA and other diffusion tensor metrics. Leveraging these models to perform streamline tractography and tractometry could provide further validation of our findings with higher spatial specificity (Kruper et al., 2021; Neher et al., 2023). Nonetheless, our study is well-situated within the large body of work on aging, white matter integrity, and physical activity using tensor-based metrics (see Maleki et al., 2022 for review). In addition, we have shown that the relationship between age and white matter integrity is *bilinear* in nature. However, the mediation models are based on *linear* relationships. As such, the age-related mediation models reported here likely underestimate the true effects. To our knowledge, no methods exist to include bilinear relationships in mediation models, and so we acknowledge the limitations of the current approach.

In line with cascade models of neurocognitive aging (e.g., Kong et al., 2019), a logical next step is exploration of how fitness and cerebrovascular health’s impact on white matter integrity may be associated with changes in cognitive outcomes, as some have done with white matter macrostructural data (Bowie et al., submitted; Breteler et al., 1994; Mace et al., 2021; Song et al., 2020; Tan et al., 2019). It would also be beneficial to leverage the regional, rather than global, measures of cerebrovascular health using Pulse-DOT to identify if vascular issues in certain brain areas are more strongly associated with aging or fitness effects than in other areas, and also to quantify arterial health indices in watershed regions, which are particularly sensitive to vascular insult (Momjian-Mayor & Baron, 2005; Torvik, 1984), and how these may impact white matter integrity.

### 4.2 Conclusions

In sum, we have demonstrated that age and fitness have non-overlapping effects on particular regions of the white matter. Aging was strongly related to loss of white matter integrity in midline regions, such as the fornix and corpus callosum (Midline FA Factor 1), but this relationship was bilinear in nature with greater loss of tissue integrity after age 64. This relationship interacted with fitness, however, such that higher fit individuals, after age 55, showed slower rate of white matter decline. Fitness, on the other hand, was uniquely related to preserved white matter integrity in deep regions that connect the brainstem and cerebellum to the thalamus and cortex (Hindbrain FA Factor 3), regardless of participant age. These relationships bore out similarly in females and males, with males exhibiting somewhat steeper age-related declines. Cerebrovascular health mediated both relationships between age or fitness and white matter integrity, indicating that the effects of age and the effects of fitness are partially explained by arterial health status. This finding underscores the importance of studying the brain within the context of the body, particularly in aging samples, in which vascular effects are known to occur and supports a cascade model of neurocognitive aging.

## Data/code availability statement

Code for analyses will be made publicly available via GitHub upon publication. Data will be made available at the end of the overarching project.

## Acknowledgments

This work was supported by NIA grants R01AG059878 and RF1AG062666 to M. Fabiani and G. Gratton. An early version of this work was presented at the 2023 meeting of the Society for Psychophysiological Research (SPR). The authors would like to thank Hannah Jones, Samantha Rubenstein, and Jeffery Gustafson for their tremendous efforts collecting data.

## Footnotes

1 Study 1: NIA grant R01AG059878; Study 2: NIA grant RF1AG062666

2 In the subsample of participants in our study who have both VO2max and eCRF, the correlation is: r(62) = 0.687, p > 0.0001, which is in line with the published literature.

3 In three cases, preprocessing failed for participants whose MPRAGE images included their entire necks as well as their heads. To remedy this, the images were snipped down to include only the head using robustfov function in FSLeyes and the preprocessing workflow was re-run.

4 tbss_1_preproc *.nii.gz / tbss_2_reg -T / tbss_3_postreg -S / tbss_4_prestats 0.2

5 Cerebral arterial elasticity measures were compared in age-matched participants from Study 1 and Study 2 using Welch’s two sample unequal variance t-tests. All metrics were not significantly different between the two studies, PTT (t(25.93) = 1.732, p = 0.095), PreFx (t(24.52) = 1.065, p = 0.2974).

6 The fourth factor’s eigenvalue was equal to 1 which violates the Kaiser rule, which states that factors less than or equal to 1 should not be retained for further analysis.

