## Supplemental Materials for "Effects of Aging, Fitness, and Cerebrovascular Status on White Matter Microstructural Health"

### Supplementary Materials

***Table S1***. *Spearman's rho correlations between PreFx and the dependent variables. All correlations are corrected for head motion and sex (except for the correlations WITH sex).*

| | Age | Fitness | Age x Fitness Interaction | Sex | Midline FA Factor 1 | Inferior-Superior FA Factor 2 | Hindbrain FA Factor 3 |
|---|---|---|---|---|---|---|---|
| PReFx | -0.42** | 0.30** | -0.04 | -0.14 | 0.17 | 0.01 | 0.11 |

***Table S2***. *Spearman's rho correlations between FA in each ROI from the JHU atlas.*

| | Age | Fitness | Age x Fitness Interaction | PTT |
|---|---|---|---|---|
| Corpus Callosum – Genu | -0.39 | 0.24 | 0.17 | 0.32 |
| Corpus Callosum – Body | -0.44 | 0.15 | 0.08 | 0.37 |
| Corpus Callosum – Splenium | -0.20 | 0.23 | 0.20 | 0.34 |
| Fornix | -0.45 | 0.05 | 0.10 | 0.30 |
| Middle Cerebellar Peduncle | -0.12 | 0.11 | -0.03 | 0.15 |
| Pontine Crossing Tract | -0.16 | 0.08 | -0.02 | 0.15 |
| Corticospinal Tract | -0.02 | 0.24 | 0.04 | 0.16 |
| Medial Lemniscus | 0.02 | 0.21 | 0.19 | 0.16 |
| Inferior Cerebellar Peduncle | -0.28 | 0.20 | 0.01 | 0.25 |
| Superior Cerebellar Peduncle | 0.31 | 0.15 | 0.08 | 0.15 |
| Cerebral Peduncle | -0.20 | 0.22 | 0.13 | 0.21 |
| Anterior Limb – Internal Capsule | -0.15 | 0.21 | 0.08 | 0.18 |
| Posterior Limb – Internal Capsule | -0.12 | 0.17 | 0.08 | 0.05 |
| Retrolenticular – Internal Capsule | -0.13 | 0.09 | 0.11 | 0.06 |

| | | | | |
|---|---|---|---|---|
| Anterior Corona Radiata | -0.42 | 0.21 | 0.17 | 0.26 |
| Posterior Corona Radiata | -0.07 | 0.09 | 0.04 | 0.12 |
| Superior Corona Radiata | -0.21 | 0.10 | 0.16 | 0.04 |
| Posterior Thalamic Radiation | -0.42 | 0.18 | 0.17 | 0.26 |
| Sagittal Stratum | -0.22 | 0.19 | 0.06 | 0.20 |
| External Capsule | -0.38 | 0.20 | 0.10 | 0.29 |
| Cingulum – Cingulate Gyrus | -0.20 | 0.18 | 0.17 | 0.23 |
| Cingulum – Hippocampus | -0.03 | 0.11 | -0.02 | 0.11 |
| Fornix (cres) / Stria Terminalis | -0.39 | 0.20 | 0.09 | 0.40 |
| Superior Longitudinal Fasciculus | -0.14 | 0.12 | 0.23 | 0.13 |
| Superior Fronto-occipital Fasciculus | -0.06 | 0.18 | 0.13 | 0.10 |
| Uncinate Fasciculus | -0.11 | 0.11 | 0.08 | 0.23 |
| Tapetum | -0.18 | -0.08 | 0.10 | 0.20 |

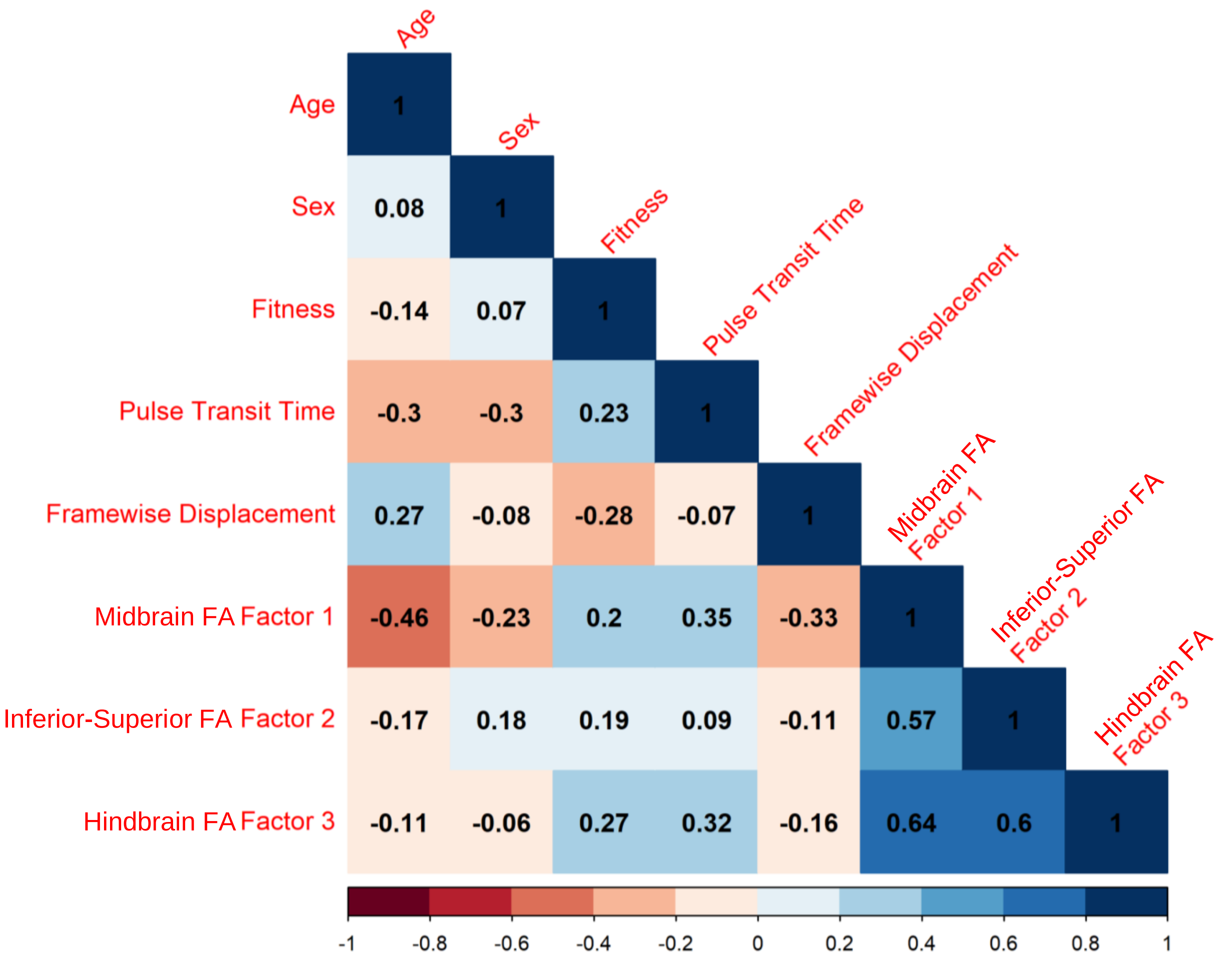


***Figure S1.*** *Spearman "raw", uncorrected correlation matrix between all the variables of interest (uncorrected for head motion or sex). Colors represent the strength of the correlations, with reddish hues indicating negative correlations and bluish hues representing positive correlations.*

**Mediation Models *without* covariates (head motion and sex) included**

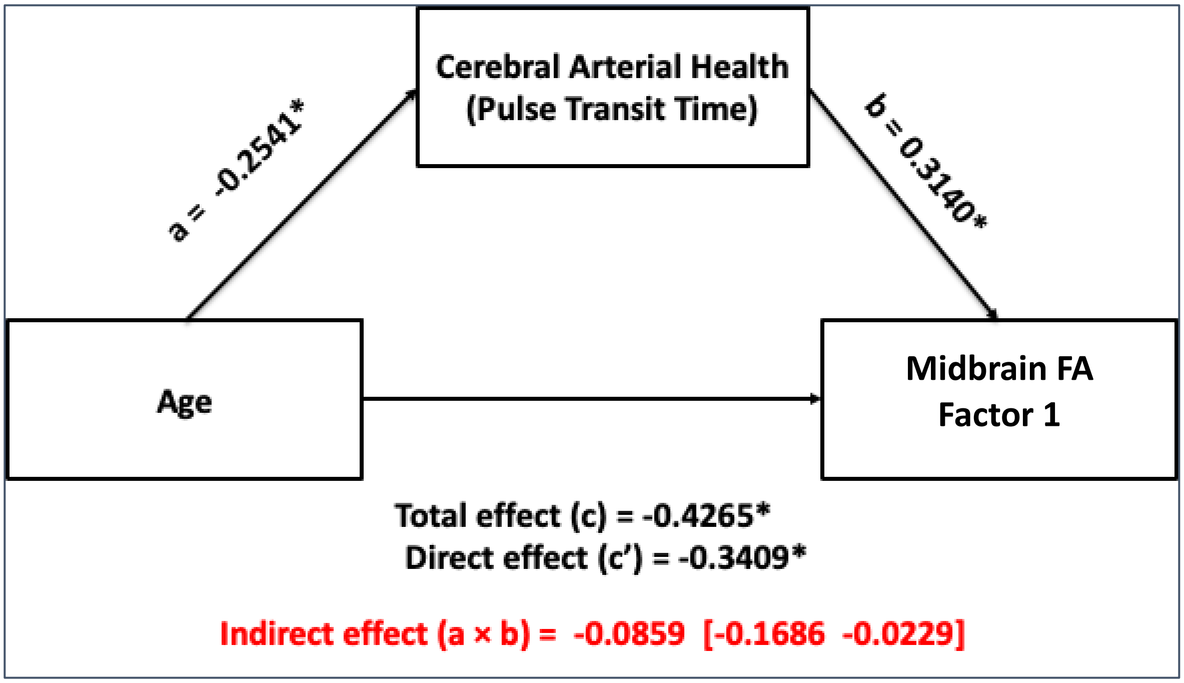


***Figure S2***. *Simple mediation between age and Midbrain FA Factor 1 using pulse transit time as a mediator without head motion or sex as covariates.*

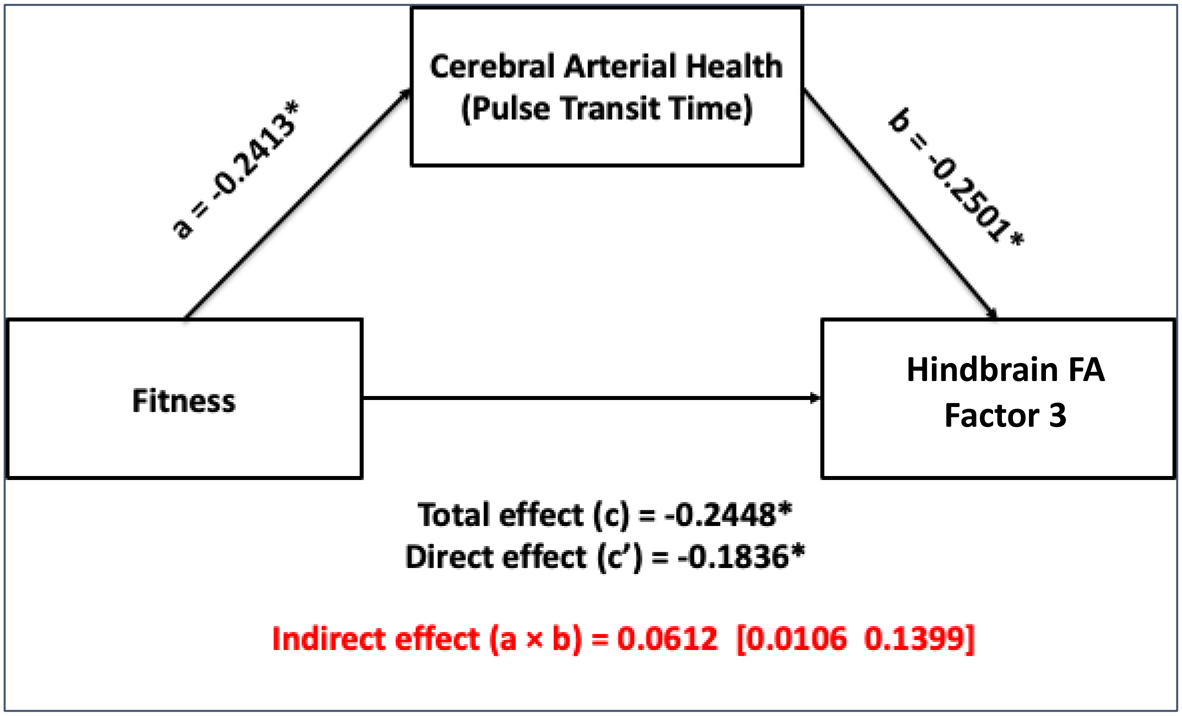


***Figure S3.*** *Simple mediation between fitness and Hindbrain FA Factor 3 using pulse transit time as a mediator without head motion or sex as covariates.*

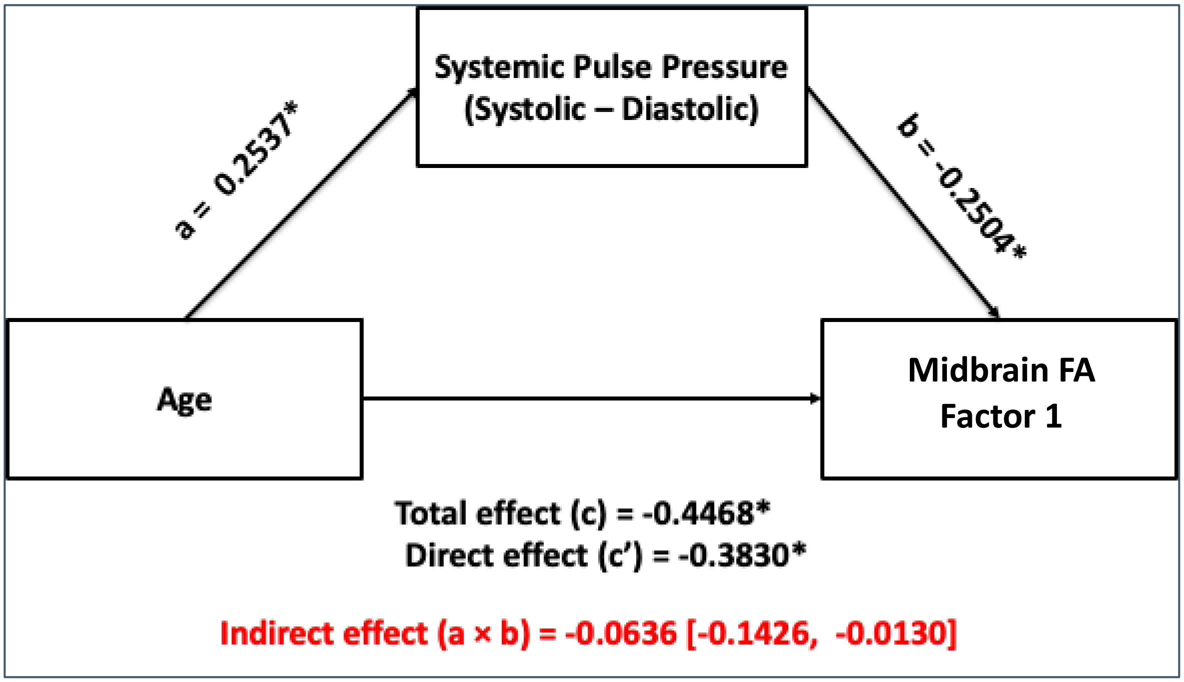


***Figure S4***. *Simple mediation between age and Midbrain FA Factor 1 using pulse pressure as a mediator without head motion or sex as covariates.*

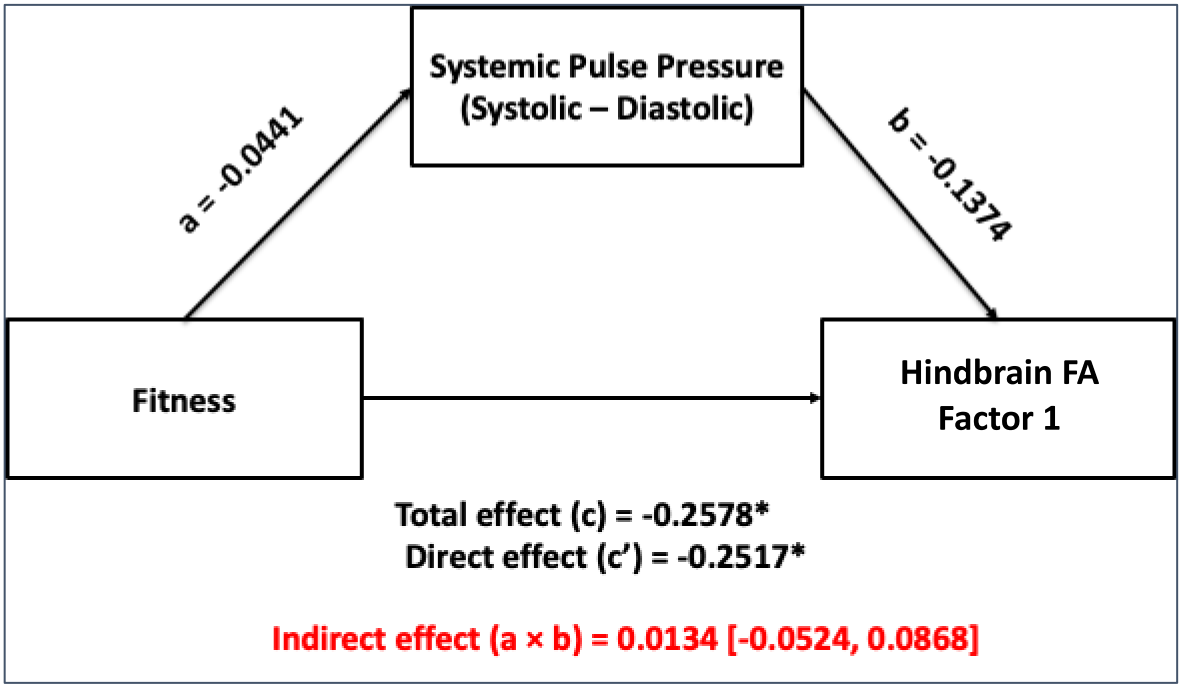


***Figure S5***. *Simple mediation between fitness and Hindbrain FA Factor 3 using pulse pressure as a mediator without head motion or sex as covariates.*

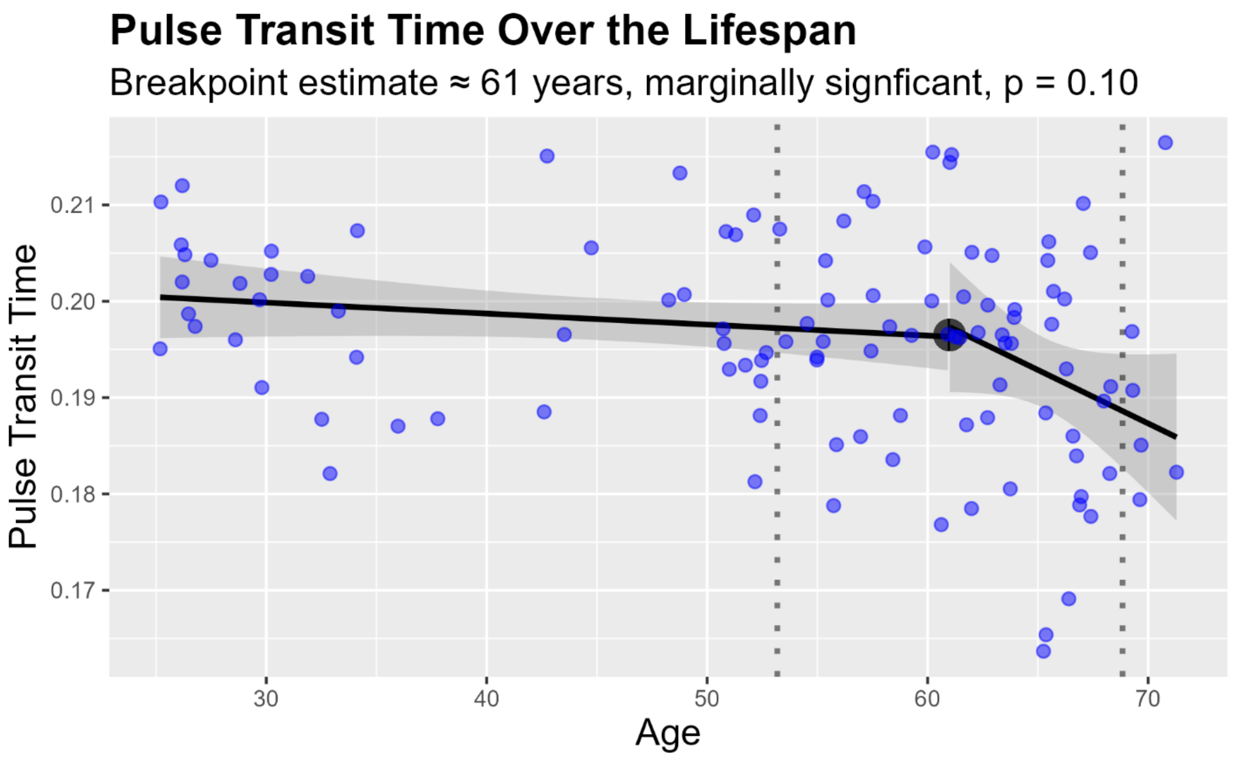


***Figure S6***. *Bilinear analysis of pulse transit time with age. The breakpoint of 61 was not significant.*

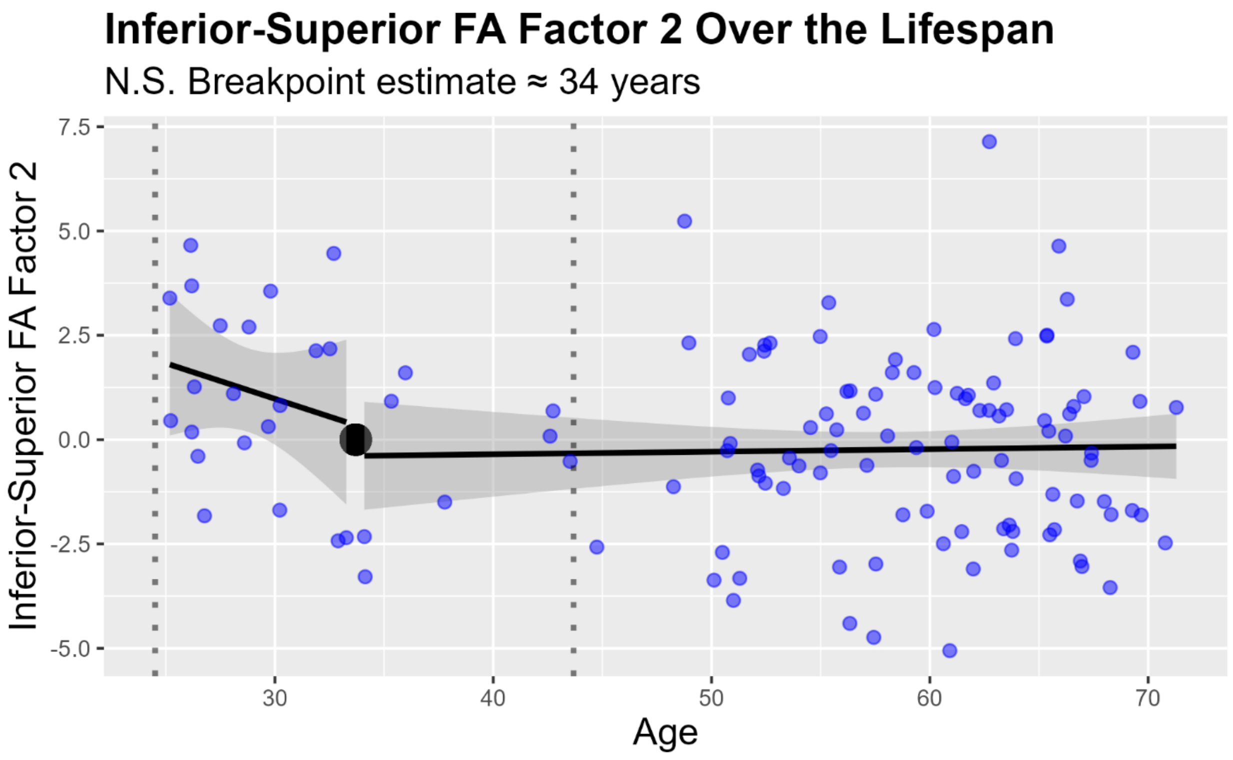


***Figure S7***. *Bilinear analysis of Factor 2 with Age. The breakpoint at 34 years was not significant.*

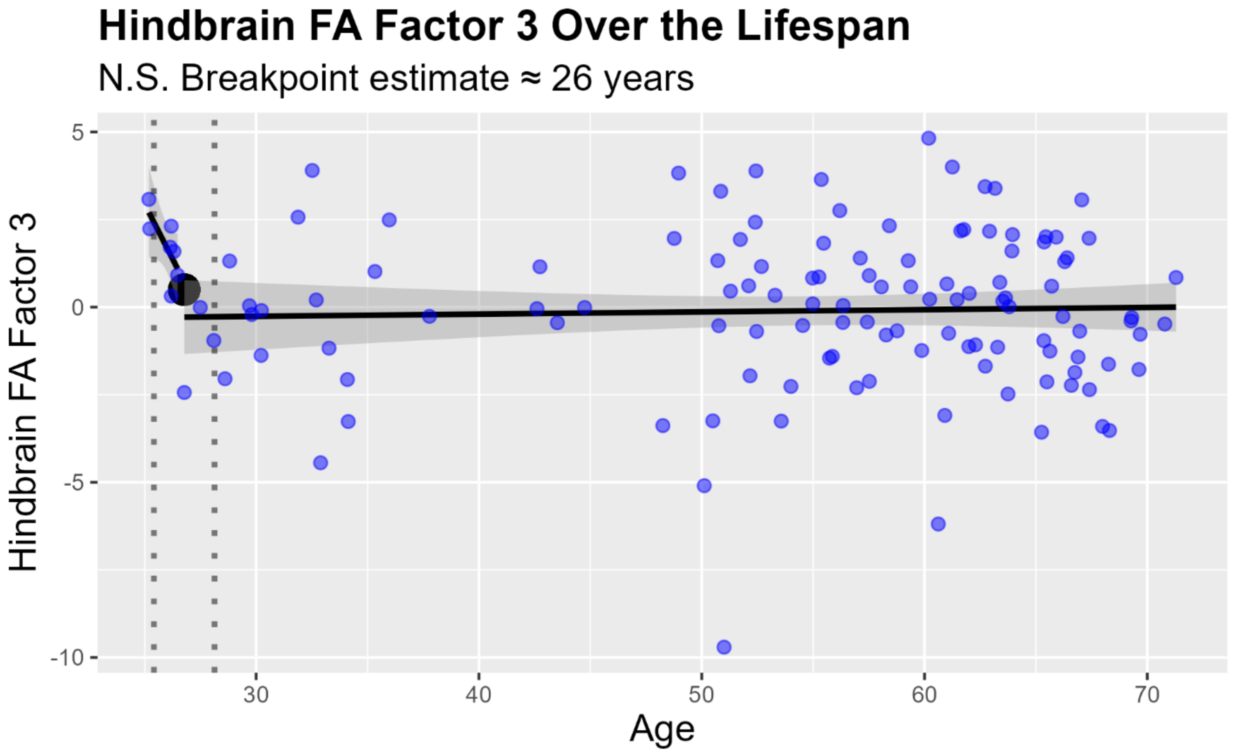


***Figure S8****. Bilinear analysis of Factor 3 with Age. The breakpoint at 26 years was not significant.*

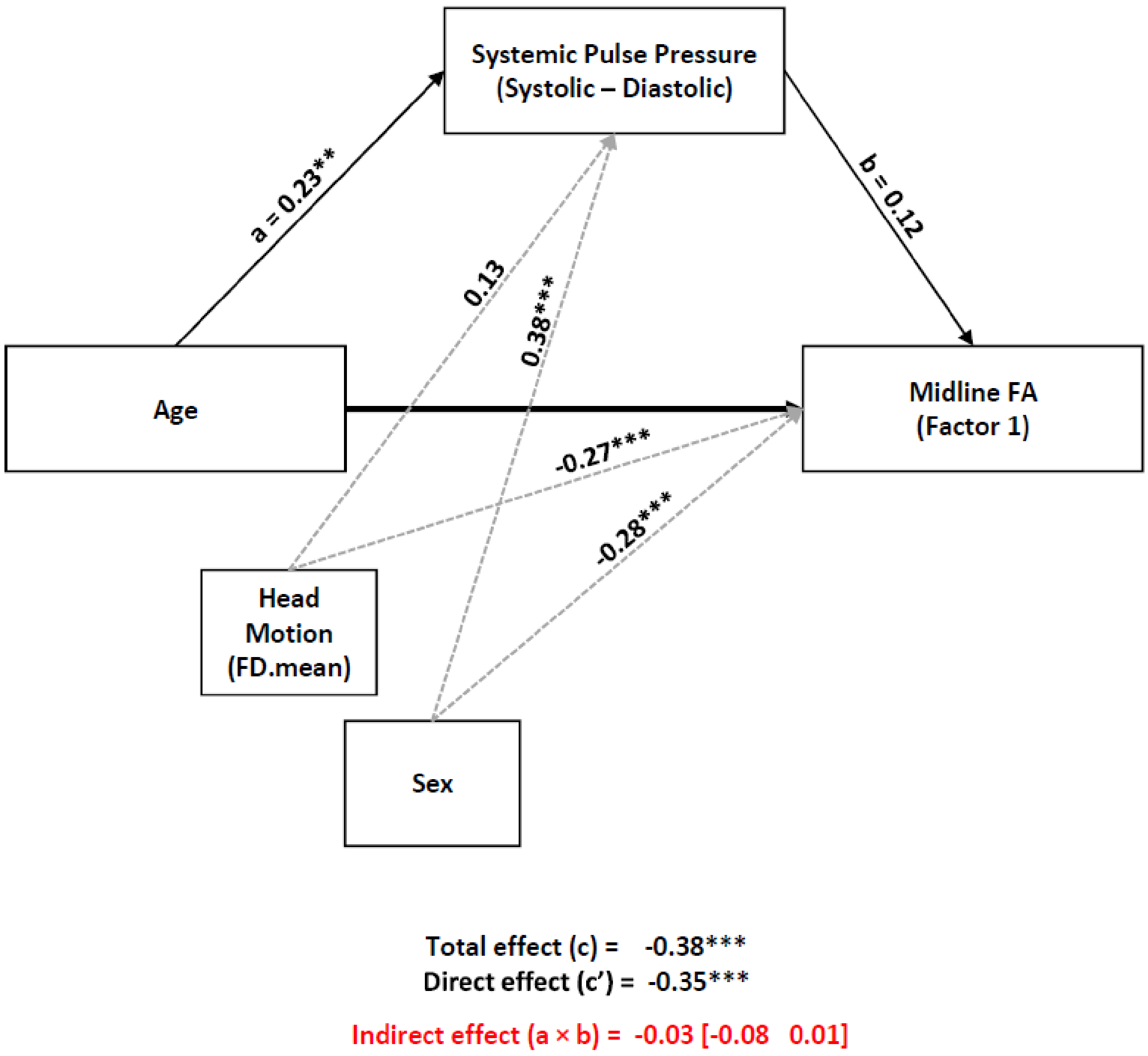


***Figure S9****. Mediation model of Age on Midline FA Factor 1 via Systemic Pulse Pressure. Head motion and Sex are covariates, as indicated by the dashed gray lines. The indirect effect is not significant. Standardized coefficients are reported. *p > .05, **p > .01, ***p > .001*

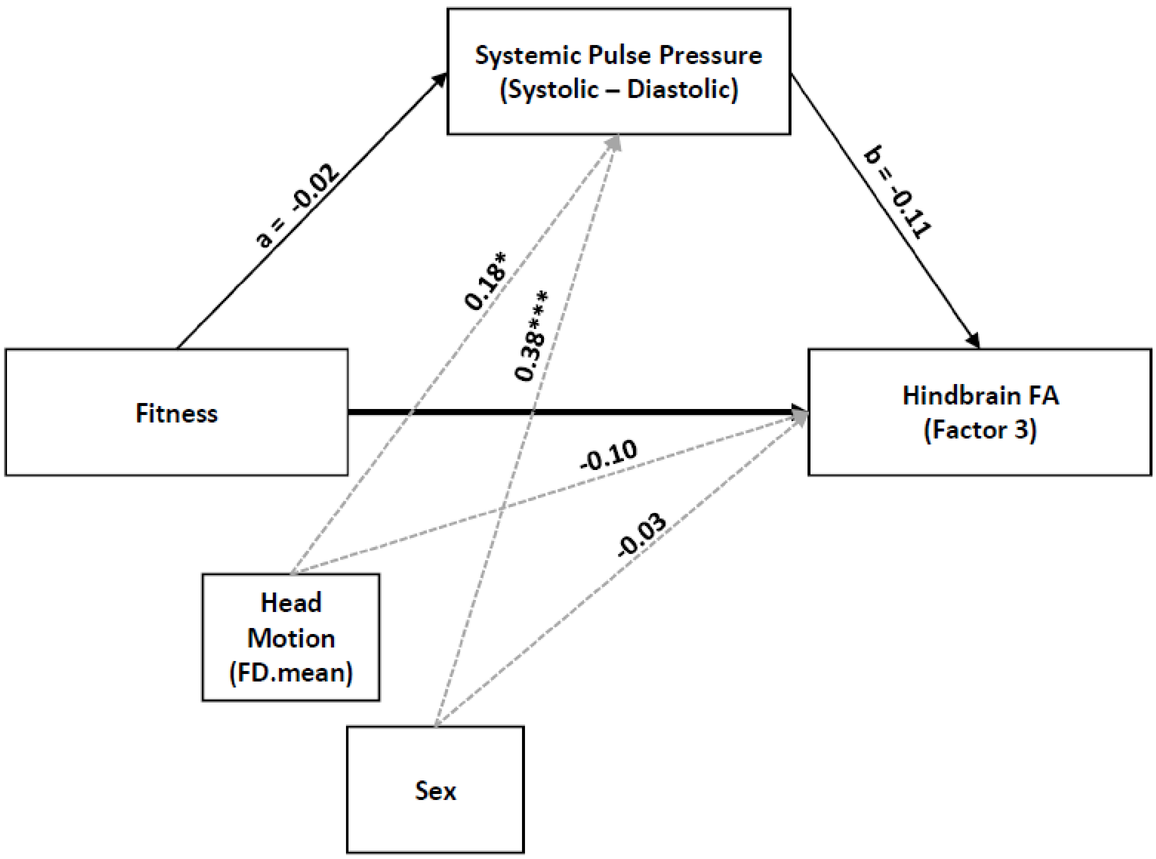


Total effect (c) = 0.228*
Direct effect (c') = 0.227*

Indirect effect (a × b) = 0.002 [-0.03 0.03]

***Figure S10***. *Mediation model of Fitness on Hindbrain FA Factor 3 via Systemic Pulse Pressure. Head motion and Sex are covariates, as indicated by the dashed gray lines. The indirect effect is not significant. Standardized coefficients are reported.* *$p < .05$, **$p < .01$, ***$p < .001$
